# Integrating tumor and immune cell transcriptomics to predict immune checkpoint inhibitor primary resistance in metastatic cutaneous melanoma

**DOI:** 10.1101/2025.08.13.670062

**Authors:** Juan Luis Onieva, Elisabeth Pérez-Ruiz, Ville Vilkki, Miguel Berciano-Guerrero, Laura Figueroa-Ortiz, Manuel Zalabardo, Beatriz Martínez-Gálvez, Isabel Barragán, Antonio Rueda-Domínguez

**Affiliations:** Group of Translational Research in Cancer Immunotherapy and Epigenetics (B-05), Medical Oncology Unit of Virgen de la Victoria Hospital, Instituto de Investigación Biomédica de Málaga y Plataforma en Nanomedicina–IBIMA Plataforma Bionand, 29010, Malaga, Spain; Facultad de Medicina, Campus de Teatinos s/n, Universidad de Málaga, 29071 Malaga, Spain; Group of Translational Research in Cancer Immunotherapy and Epigenetics (B-05), Medical Oncology Unit of Regional Hospital, Instituto de Investigación Biomédica de Málaga y Plataforma en Nanomedicina–IBIMA Plataforma Bionand, 29010, Malaga, Spain; Group New Horizons for Cancer Patients (B-23), Medical Oncology Unit of Regional Hospital, Instituto de Investigación Biomédica de Málaga y Plataforma en Nanomedicina–IBIMA Plataforma Bionand, 29010, Malaga, Spain; Group of Cancer Clinical and Translation Research (B-01), Medical Oncology Unit of Virgen de la Victoria Hospital, Instituto de Investigación Biomédica de Málaga y Plataforma en Nanomedicina–IBIMA Plataforma Bionand, University of Malaga, 29010, Malaga, Spain; Group of Pharmacoepigenetics, Department of Physiology and Pharmacology, Karolinska Institutet, 171 65, Stockholm, Sweden

**Keywords:** Melanoma, Predictive markers, Prognostic markers, Omics integration, Immune Checkpoint Inhibitors, Treatment response, Primary resistance

## Abstract

**Background:** The emergence of immune checkpoint inhibitors (ICIs) has transformed the treatment landscape of metastatic melanoma. However, despite its success, reliable biomarkers for predicting primary resistance are not available in clinical practice. This study seeks to identify predictors of primary resistance based on novel gene expression signatures using pre-treatment multidimensional profiling in melanoma patients.

**Methods:** The transcriptomic profile of the tumor microenvironment was analyzed using tissue samples from 46 metastatic cutaneous melanoma patients collected prior to the initiation of ICIs therapy. A primary resistance predictive model was trained with the Discovery FFPE RNA-seq sub-cohort and validated using an independent external cohort of 54 samples. Additionally, liquid biopsy samples from peripheral blood mononuclear cells were analyzed in 8 patients using single-cell RNA sequencing (scRNA-seq) and in 46 patients using flow cytometry to characterize the distribution and abundance of the different immune cell populations.

**Results:** We identified an 82-gene transcriptomic signature composed of tumor- and immune-related genes that stratifies metastatic cutaneous melanoma patients based on primary resistance to ICIs, with key markers including *CXCL13, WDR63, MZB1, FDCSP, IGKC* and *GRIK3*. This signature was enriched for pathways related to B cell activation and immune cell communication and achieved an AUC of 0.814 in predictive modeling. Immune deconvolution guided by scRNA-seq revealed four immune cell subsets (Plasma cells, Pre-B cells, Memory CD4⁺ T cells, and Naive CD4⁺ T cells) as prognostic indicators of resistance. Some of these subpopulations were validated by flow cytometry before and after treatment.

**Conclusions:** We propose a transcriptomic biomarker signature that accurately predicts primary resistance to ICIs in metastatic cutaneous melanoma. Through the integration of immune deconvolution with circulating immune cell profiles, we derived an ImmuneSignature linked to patient survival. By combining these approaches, we provide a framework for enhancing the prediction of immunotherapy outcomes and offer a novel strategy for identifying therapeutic targets to overcome resistance. Our findings lead to more effective and personalized immunotherapy guidance.

## Background

The development of immune checkpoint inhibitors (ICIs) therapies has dramatically changed cancer treatment paradigm, providing substantial clinical benefits for patients with previously hard-to-treat cancers, including metastatic melanoma and non-small cell lung cancer among other solid tumors (1,2). ICI therapies target immune checkpoint proteins, such as PD-1, PD-L1, and CTLA-4, lifting inhibitory signals on T cells and allowing them to reactivate their cytotoxic function to identify cancer cells (3). However, despite its promising potential, response rates are highly variable. Although some patients achieve prolonged tumor control, nearly 60-70% (4) experience either primary or acquired resistance to therapy (5). This variability in treatment efficacy emphasizes a critical need for reliable biomarkers to predict patient responses, optimize patient selection, and reduce treatment costs.

Currently, PD-L1 protein expression, microsatellite instability (MSI), and tumor mutational burden (TMB) are considered standard of care biomarkers, recognized by FDA and EMA, for predicting response to ICIs therapy in specific clinical settings (4). However, these factors alone do not fully account for the complexity of intratumor heterogeneity, limiting their predictive accuracy (6). To address these limitations, gene expression profiling has been established as a solid strategy, offering a detailed view of the tumor-immune microenvironment (TIME) and facilitating the identification of molecular signatures linked to ICIs response in other cancer types (7,8).

The identification of patients likely to exhibit primary resistance remains a critical challenge for the optimized use of ICIs, preventing prolonged treatment without clinical benefit for the patients. To date, numerous transcriptomic biomarkers and computational methods have been proposed to predict responses to ICI therapies in metastatic melanoma (7,9–14). However, these approaches face several challenges, including a lack of standardized response criteria (4). To overcome this limitation, the Society for Immunotherapy of Cancer (SITC) established a consensus framework for the evaluation of primary resistance to ICIs (5). In this study, we analyzed transcriptomic data from patients with metastatic cutaneous melanoma treated with anti-PD-1 therapy to identify a gene signature that predicts primary resistance by SITC criteria. Using an integrated approach with bulk and single-cell transcriptomic data, from tissue and liquid biopsy respectively, we leveraged cell-specific gene expression profiles to enhance the precision of tissue deconvolution. This approach allows for more accurate estimation of immune landscape, capturing immune cell populations that are mirrored in peripheral blood composition (15,16). To further validate our findings and enhance clinical relevance, we performed an independent flow cytometry analysis of peripheral blood mononuclear cells (PBMCs) from patients with metastatic cutaneous melanoma in the same treatment context. This additional validation step not only enhances the robustness of our results but also provides insights into the potential applicability of ICI resistance biomarkers in non-invasive, peripheral blood-based assays.

In this regard, the goal of the study was to identify a molecular signature linked to primary resistance to ICIs in metastatic cutaneous melanoma. Furthermore, the study aimed to formulate a predictive model while highlighting immune cell subsets that may serve as prospective therapeutic targets.

## Methods

### Cohorts of melanoma patients

The patients included in this study are metastatic cutaneous melanoma patients that received ICIs in monotherapy (anti-PD1) or in combination with tyrosine kinase inhibitors. The Discovery cohort includes 89 patients recruited between 2015 and 2023, across two centers: Hospital Universitario Regional de Málaga and Hospital Universitario Virgen de la Victoria. Solid and liquid samples from these patients were processed for multidimensional evaluation that led to the primary resistance prediction model and the prognostic ImmuneSignature. Each of the dimensions defines the three subdivisions of the Discovery cohort: Formalin-Fixed Paraffin-Embedded (FFPE) bulk RNA-seq (Discovery FFPE bulk-RNA seq sub-cohort, 46 samples, pre-treatment); PBMCs scRNA-seq (Discovery Single-Cell sub-cohort, 8 samples, pre-treatment) and Flow cytometry (Discovery Flow cytometry sub-cohort, 46 samples, before ICI initiation (T1) and after the second cycle of ICIs –2 or 3 weeks–(T2)). Patient overlapping between any of the 2 sub-cohorts was 8 at maximum and 1 patient between the 3 sub-cohorts.

For the predictive and prognostic markers validations, we have used the Gene Expression Omnibus (GEO) database repository to compile 6 datasets of metastatic melanoma patients with pre-ICI treatment transcriptomic data from clinical and translational studies. Only patients with cutaneous melanoma subtype were retained. These datasets have been organized in two validation cohorts: the Validation cohort (n=54) in which progression-free survival (PFS) data is available and has been used for the independent external validation of the primary resistance model (7), and the Extended Validation cohort (n=286) where only overall survival (OS) data is available and served for the validation of the prognostic score (7,9–13).

### Inclusion and exclusion criteria

Participants were eligible for inclusion if they met the following inclusion criteria: histological confirmation of metastatic cutaneous melanoma, age ≥18 years, and normal hepatic, renal, cardiac, and bone marrow function, as determined by standard clinical laboratory tests. Additionally, for the Discovery cohort, the availability of a pre-treatment biopsy with sufficient tumor material for histological and molecular analyses was required. Enrollment was independent of the cancer stage or personal and family history of cancer at the time of diagnosis. Exclusion criteria were refusal to participate in the study after providing informed consent, contraindications to receiving immunotherapy, or the presence of uncontrolled infectious diseases. Patients with coexisting neoplastic diseases other than metastatic melanoma or with autoimmune diseases were also excluded.

All data were extracted from complete and existing medical records, resulting in no attrition for loss of follow-up.

### Definition of primary resistance

Throughout the multidimensional analysis, patients were categorized into non-resistant and resistant patients, defined by primary resistance (5): non-resistant patients are characterized by RECIST v1.1-calculated tumor response or prolonged stable disease lasting ≥ 6 months; resistant patients experienced disease progression before 6 months after receiving at least two complete cycles of ICIs.

### Ethics

Patients sample collection adhered to the Declaration of Helsinki and was approved by the Comité de Ética de la Investigación Provincial de Málaga (Approval: 26 October 2017, Project Title: “Omics integration for precision cancer immunotherapy,” 799818, H2020-MSCA-IF-2017). All patients provided informed consent. Participant data were collected from February 5, 2015, to May 29, 2023, across two centers: Hospital Universitario Regional de Málaga and Hospital Universitario Virgen de la Victoria.

### Bulk and single-cell RNA profiling

RNA from the 46 tumor samples of the Discovery FFPE bulk RNA-seq sub-cohort was extracted with the RNeasy FFPE kit following the manufacturer’s instructions (Qiagen, Düsseldorf, Germany; Ref. 73504) and RNA-Seq libraries were prepared using the TruSeq Stranded Total RNA Gold kit (Illumina, Ref. 20020598) with IDT for Illumina TruSeq RNA UD Indexes (Illumina, Ref. 20020591), capturing both coding and non-coding RNA through double ribosomal RNA depletion. Sequencing coverage and quality statistics for bulk RNA-seq are provided in supplementary table S7.

Single-cell suspensions of cryopreserved PBMCs from 15 mL of blood of 8 patients (Discovery Single-cell sub-cohort) were loaded onto the Chromium Controller to generate single-cell gel bead-in-emulsions (GEMs), capturing individual cells with barcoded mRNA. Libraries were then prepared according to the 10x Genomics Single-Cell 3’ Library and Gel Bead Kit v3.1 Dual Index protocol (CG000315 Rev E). Illumina NextSeq 550 and NovaSeq 6000 platforms were used for sequencing, and batch effects were accounted for in all analyses.

### RNA-seq raw data processing

Bulk RNA-seq fastq quality control was performed with FastQC (RRID:SCR_014583), and reads were trimmed using HISAT2 (v2.1.0) (RRID:SCR_015530) with a rRNA index. Trimmed fastq files were aligned to the GRCh38 reference genome using STAR (v2.5.1b) (RRID:SCR_004463), with read quantification and mapping quality evaluated through featureCount. Raw scRNA-seq sequencing data were aligned with Cell Ranger against the GRCh38 reference genome. Data were normalized and preprocessed with Seurat v5 (RRID:SCR_016341) (17), with cell populations visualized using PCA and UMAP and identified through sc-type algorithm (18). Louvain clustering with shared nearest neighbor modularity was applied to identify cell clusters, with additional refinement for B cell lineage resolution by adjusting community parameters.

### Multidimensional analysis and integration to predict primary resistance

Differential expression (DE) analysis in the Discovery FFPE bulk-RNA seq sub-cohort samples (n=46) was conducted using DESeq2 (RRID:SCR_015687), considering genes as DE if they had a baseMean count >10, absolute log2FC >1.5, and an adjusted p-value <0.05. For pathway analysis, Ingenuity Pathway Analysis (IPA, QIAGEN) (RRID:SCR_008653) was employed to interpret the biological significance of DE genes and identify pathways relevant to ICIs response.

7-fold cross-validation framework repeated 1000 times was the statistical technique used for internal validation in the Discovery FFPE bulk RNA-seq sub-cohort (training cohort). An Elastic Net as classification model was applied over the DE genes expression to build a predictive model for primary resistance to ICIs. The model was implemented using the caret package in R, with the glmnet function. The regularization parameter lambda and the mixing parameter alpha were optimized during internal validation using a grid search approach. The optimal hyperparameters were selected based on the highest mean AUC obtained across cross-validation folds, with lambda set to 0.19 and alpha set to 0.5. To control for overfitting, model selection and hyperparameter tuning were strictly confined to the training dataset, and final model evaluation was carried out on an independent dataset, the Validation cohort (n=54), to assess the generalizability of our proposed predictive signature. The performance of the model was evaluated using the area under the curve (AUC) as metric, with 95% confidence intervals computed via bootstrapping (n = 1000 resamples). Variable importance was extracted using the varImp function from the caret package, which computes importance scores based on the magnitude of the model coefficients derived from the fitted glmnet model. Given the moderate imbalance (41% resistant vs. 59% non-resistant), no explicit correction was applied to address class imbalance in the training set. Subgroup analyses were not performed due to the limited sample size. However, this limitation is partially mitigated by the design of a clinically homogeneous cohort: all patients received anti-PD-1 therapy and shared the same histological subtype—cutaneous melanoma—minimizing clinical heterogeneity that could otherwise confound model performance.

In the Discovery Single-Cell sub-cohort samples (n=8), marker genes within each cluster were identified using Seurat’s FindMarkers function with MAST test, which highlighted genes with significant differential expression across cell types and primary resistance. Sex and batch were included as covariates in the design formula for the DE analysis to account for potential confounding effects in gene expression.

To assess the contribution of each cell type to the bulk RNA-seq signal with higher resolution and specificity for metastatic cutaneous melanoma, non-matrix factorization was applied using the identified marker genes from the scRNA-seq analysis (15). This method provided a quantitative estimation of the relative abundance of the different circulating immune cell populations within the tumor sample.

### Flow Cytometry

PBMCs, isolated from 10 mL of blood collected at pre-treatment (T1) and post-treatment (T2), were obtained from the Discovery Flow Cytometry sub-cohort (46 patients) and characterized by flow cytometry using a BD FACS Canto II cytometer (BD Biosciences, San Jose, CA, USA). Two different panels were used for the staining. In the first panel, T cells were identified using FITC anti-human CD4 antibody (BioLegend, Cat# 317408). In the second panel, Plasma cells were identified using the following monoclonal antibodies: PE anti-human CD20 (BioLegend, Cat# 302306), FITC anti-human CD138 (Syndecan-1, BioLegend, Cat# 356508), and PE/Cyanine7 anti-human CD27 (BioLegend, Cat# 302838). Initial gating was performed on forward scatter and side scatter to control event size and granularity properties. T cells were gated as CD4^+^ events within the lymphocyte gate. Plasma cells were defined as CD27^+^ CD138^+^ events within the CD20^-^ population. Data from the two time points were acquired using DIVA software (v9.0) and subsequently downloaded for detailed analysis with flowTOTAL (19).

### Survival analysis and ImmuneSignature

Survival analysis was performed with the Kaplan–Meier method to estimate the survival curves. The log-rank test, implemented in the R package survival, was applied to assess statistical significance in the survival differences. To evaluate the prognostic value of immune-related features, the maximally selected log-rank statistic was applied individually to each of the four immune cell populations of interest (Plasma cells, Pre-B cells, Memory CD4^+^ T cells, and Naive CD4^+^ T cells), dichotomizing patients into high and low groups based on estimated cell proportions within the tumor. The resulting classifications were then used to define a composite ImmuneSignature as follows: samples were assigned to the ImmuneSignature-High group if either (i) both Plasma cells and Pre-B cells were classified as high, or (ii) both Memory CD4^+^ T cells and Naive CD4^+^ T cells were classified as high. All remaining combinations were classified as ImmuneSignature-Low. The prognostic impact of the ImmuneSignature was subsequently assessed using log-rank tests for OS.

For flow cytometry analysis, a surrogate ImmuneSignature was constructed using the percentage of immune populations detectable with the available antibody flow cytometry panel in liquid biopsy samples. While the original signature was based on four immune cell types inferred from transcriptomic deconvolution in tumor tissue, only CD4^+^ T cells and Plasma cells were included in the flow cytometry panel. As a result, the flow-based ImmuneSignature represents a partial implementation of the original tissue-based ImmuneSignature. This reflects a key technical limitation of the study, as the reduced marker coverage may limit the ability to fully recapitulate the tissue-derived ImmuneSignature in peripheral blood.

### Statistical analysis

All statistical analyses were conducted using R (version 4.2.0; RRID:SCR_001905) and relevant CRAN and Bioconductor packages. Continuous variables were summarized using medians, while categorical variables were presented as counts and percentages. Survival analyses were performed using Kaplan–Meier estimates, with differences between groups assessed via the log-rank test. The Wilcoxon test was used to compare continuous variables between non-resistant and resistant patients. An overview of the methodological workflow is provided in figure 1.

**Figure 1.**
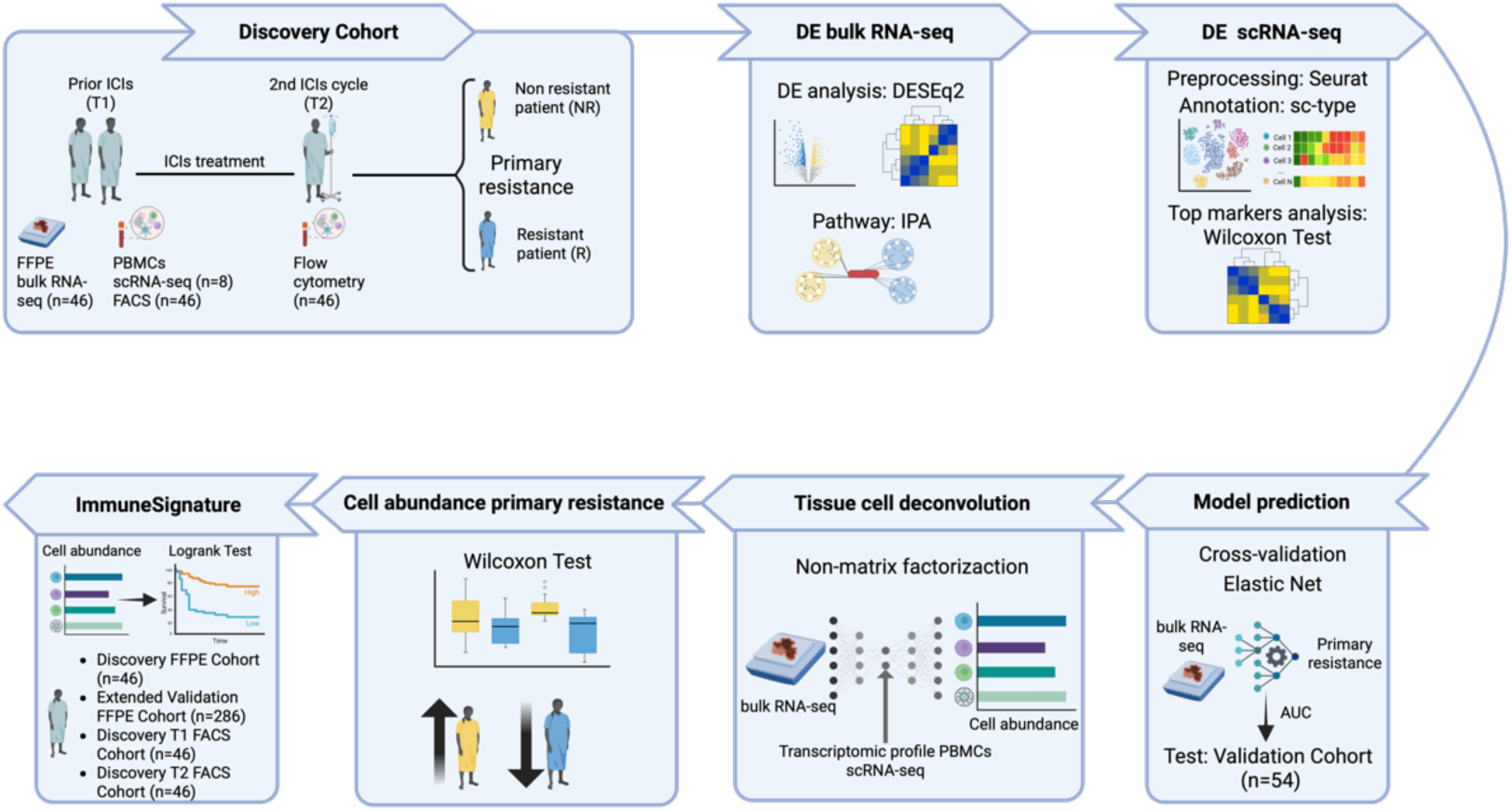
Schematic overview of the methodological workflow employed in this study. The diagram outlines the sequential steps from data acquisition and preprocessing, through analysis and model development, to validation.

## Results

In total, 89 patients with metastatic cutaneous melanoma were collected for the Discovery cohort of this study. Baseline characteristics of the patients are summarized in the supplementary table S1. Overall, 42 (47.19%) patients were female, median age was 64 years (interquartile range 50–73 years). In terms of primary resistance, 49 patients were classified as non-resistant (55.06%) and 40 (44.94%) patients as resistant.

The median PFS for patients with primary resistance was 2.29 months (95% CI: 1.60–3.24), whereas the median PFS for non-resistant patients was 28.72 months (95% CI: 17.67–NR) (supplementary figure S1A). Regarding OS, the median for resistant patients was 5.62 months (95% CI: 3.27–11.27), while non-resistant patients had a significantly longer OS, with a median of 34.26 months (95% CI: 27.05–NR) (supplementary figure S1B).

In supplementary table S1, we provide a detailed summary of the variables segregated by the sub-cohorts outlined in the Methods section—namely, the FFPE bulk-RNA seq, Single-Cell, and Flow cytometry cohorts. These cohorts share comparable baseline clinical characteristics, with no significant differences in key parameters. Patient overlapping is minimal.

### Baseline B cell-enriched transcriptomics signature in non-resistant patients to ICIs

We initially focused on the Discovery FFPE bulk-RNA seq sub-cohort, which consisted of 46 patients. Bulk RNA sequencing was conducted on samples collected prior to the initiation of immunotherapy, enabling the comparison of gene expression between patients classified as resistant (n=19) and non-resistant (n=27) based on primary resistance. Analysis of the transcriptomic profiles revealed a total of 82 DE genes (supplementary table S2). Of these, 60 genes were upregulated in non-resistant patients, while 22 genes showed higher expression in resistant patients (figure 2A). Among the downregulated genes in non-resistant patients, *GRIK3* and *LINC01502* exhibited the most significant changes. On the other hand, the upregulated genes were enriched in genes related to the immune system, such as immunoglobulin families and B cell-associated genes like *MZB1*.

**Figure 2.**
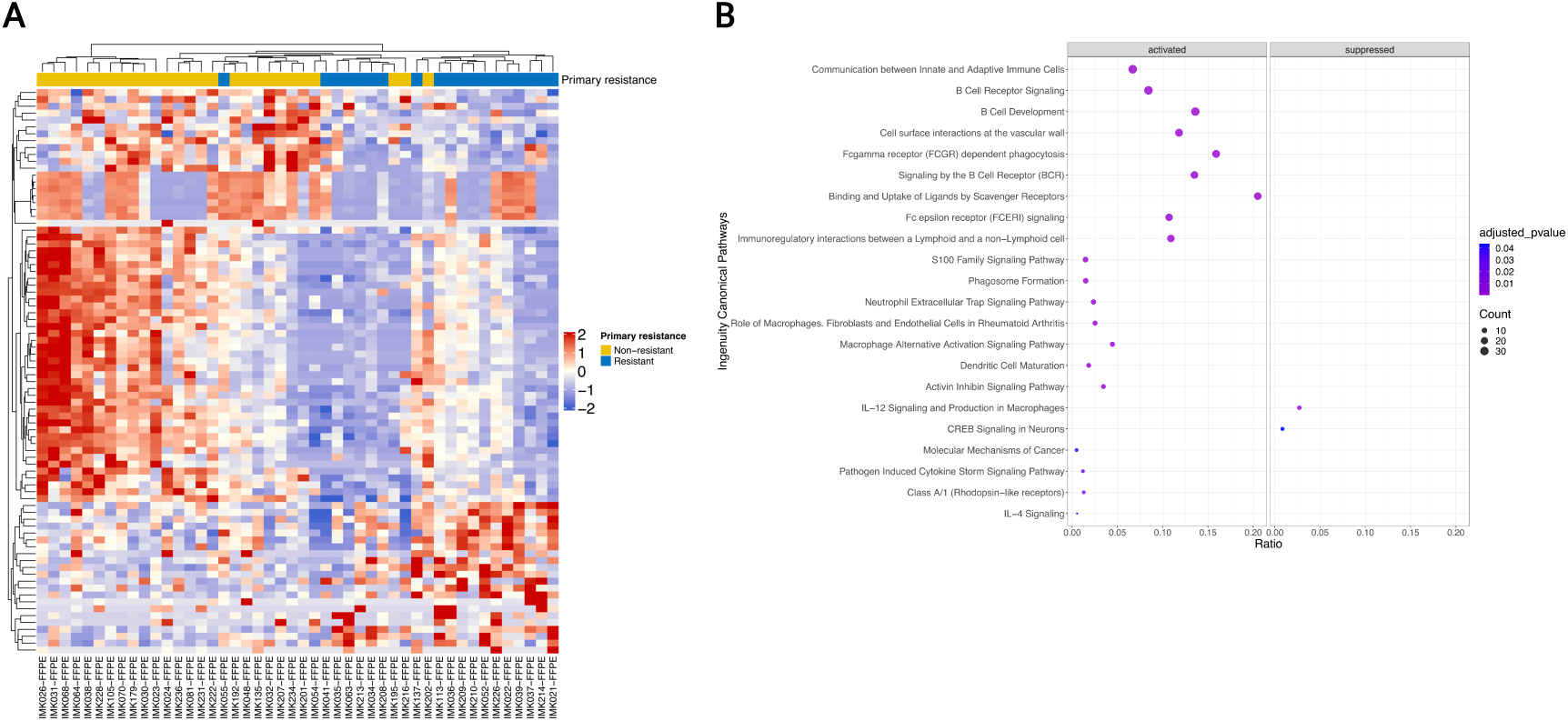
Gene expression and pathway enrichment in primary resistance analysis. (A) Heatmap depicting the differential expression of 82 genes between resistant and non-resistant patients, with color intensity representing the level of expression in each FFPE Bulk-RNA seq sample. (B) Dot plot of the Ingenuity Canonical Pathways, showing significant enrichment of pathways in non-resistant patients, categorized as activated or suppressed. The size of each dot reflects the number of DE genes related to the pathway while the color indicates the significance of the enrichment.

To further elucidate the biological pathways underlying the identified genes, pathway analysis performed in IPA revealed activation of both the innate and adaptive immune systems. The *Communication between Innate and Adaptive Immune Cells* pathway, comprising 30 upregulated genes in non-resistant patients, exhibited the most robust gene involvement and was significantly activated in non-resistant patients. Similarly, several B cell–related processes— including *B Cell Receptor Signaling*, *B Cell Development, and Signaling by the B Cell Receptor (BCR)—*were enriched and activated in non-resistant patients. In resistant patients, significant activation was observed in the *IL-12 Signaling and Production in Macrophages* and *CREB Signaling in Neurons* pathways. IL-12 signaling in macrophages modulates the immune response and promotes inflammation and angiogenesis through pathways such as NF-kB and JAK/STAT. Additionally, CREB signaling enhances cell survival, metastasis, and resistance to apoptosis through pathways like PKA, mTOR, and Notch, which are involved in tumor cell proliferation and invasion. These findings highlight opposed immunological profiles between the two groups, suggesting that enhanced immune communication and B cell activity may play critical roles in mediating primary non-resistance to immunotherapy, while angiogenesis and anti-apoptotic pathways amongst others may drive resistance mechanisms (figure 2B).

### Predictive model of primary resistance to ICIs from bulk transcriptomics of the TIME

The genes identified in the differential analysis were used to generate a predictive classification model using the Elastic Net algorithm, with non-resistant and resistant patients as the target labels. The model was trained using 7-fold cross-validation, allowing for hyperparameter tuning during internal validation using the Discovery FFPE bulk-RNA seq sub-cohort as training cohort. The AUC was employed as the performance metric to evaluate the model. The trained model was subsequently tested on the Validation cohort consisting of 54 patients (7). The resulting AUC for the model was 0.814 (95% CI: 0.701–0.923) (figure 3A), with a positive predictive value of 0.810 and a sensitivity of 0.781. The overall performance of the model accurately excelled the identification of true negatives, the resistant patients.

**Figure 3.**
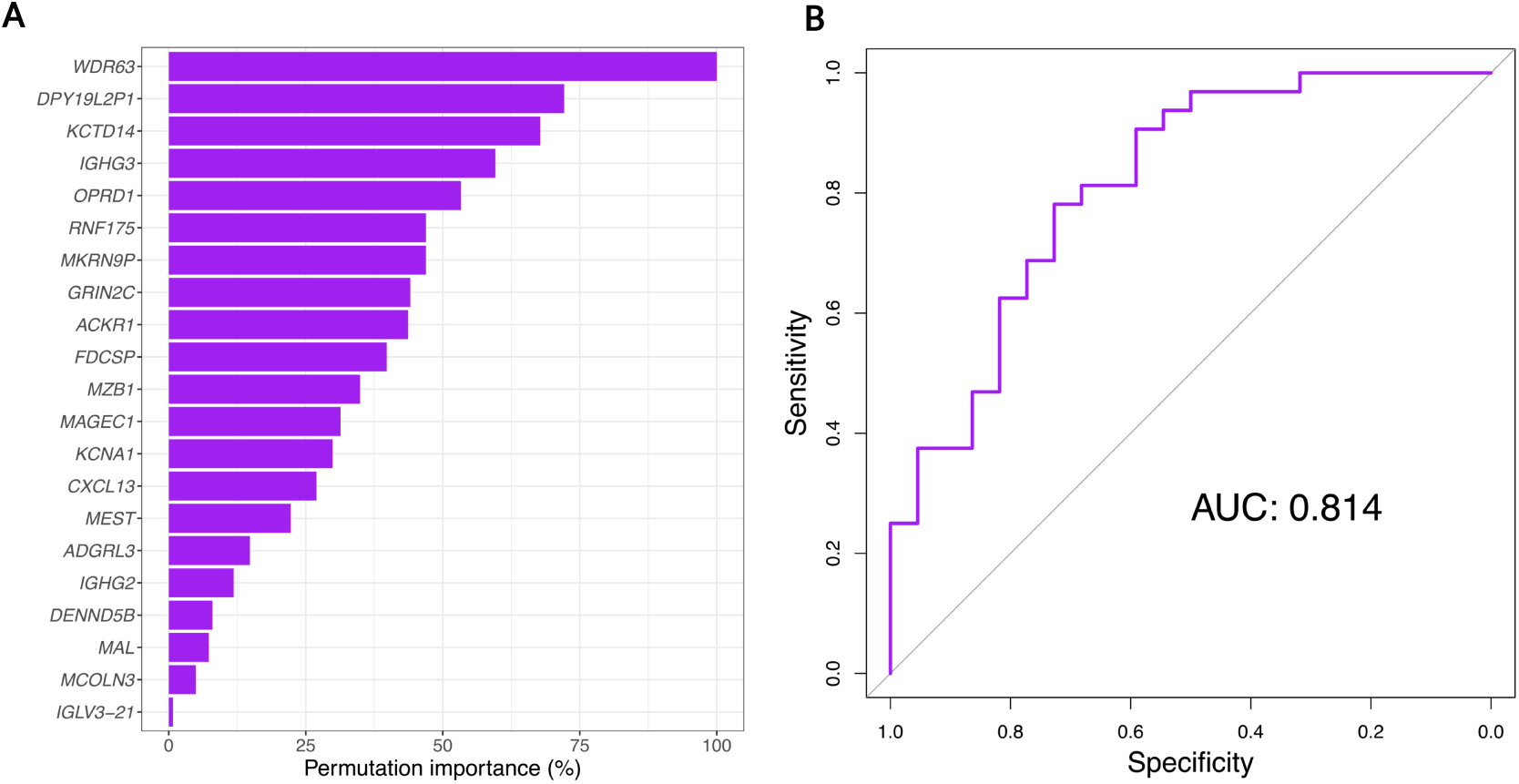
Model performance in predicting primary resistance in an independent melanoma cohort. (A) Receiver operating characteristic (ROC) curve showing the performance of the predictive model trained on the discovery cohort and tested on additional 54 melanoma samples from an independent external cohort. The model achieved an AUC of 0.814. (B) Variable importance plot showing the top features selected by the model based on permutation importance scores. Genes such as *WDR63* exhibit the highest importance in predicting the primary resistance.

The model’s predictive capabilities were characterized through a variable importance analysis, highlighting the key features contributing to its performance. Based on relevance scores, genes such as *WDR63* emerged as the most critical determinants in predicting primary resistance (figure 3B).

### Identity and primary resistance markers from single-cell transcriptomics of the PBMCs

The Discovery Single-Cell sub-cohort (n=8) was employed to phenotypically characterize the different cell types within the patients PBMC samples using their single-cell-specific gene expression profiles. We identified up to 12 distinct populations based on the top markers embedded in the sc-type algorithm (supplementary figure S2): CD8^+^ NKT-like cells, Classical monocytes, Effector CD4^+^ T cells, Effector CD8^+^ T cells, Memory CD4^+^ T cells, Naive CD4^+^ T cells, Natural killer cells, Non-classical Monocytes, Plasma B cells, Plasmacytoid dendritic cells, Platelets, and Pre-B cells. In figure 4A, the UMAP visualization shows the distinct populations identified in the scRNA-seq analysis.

**Figure 4.**
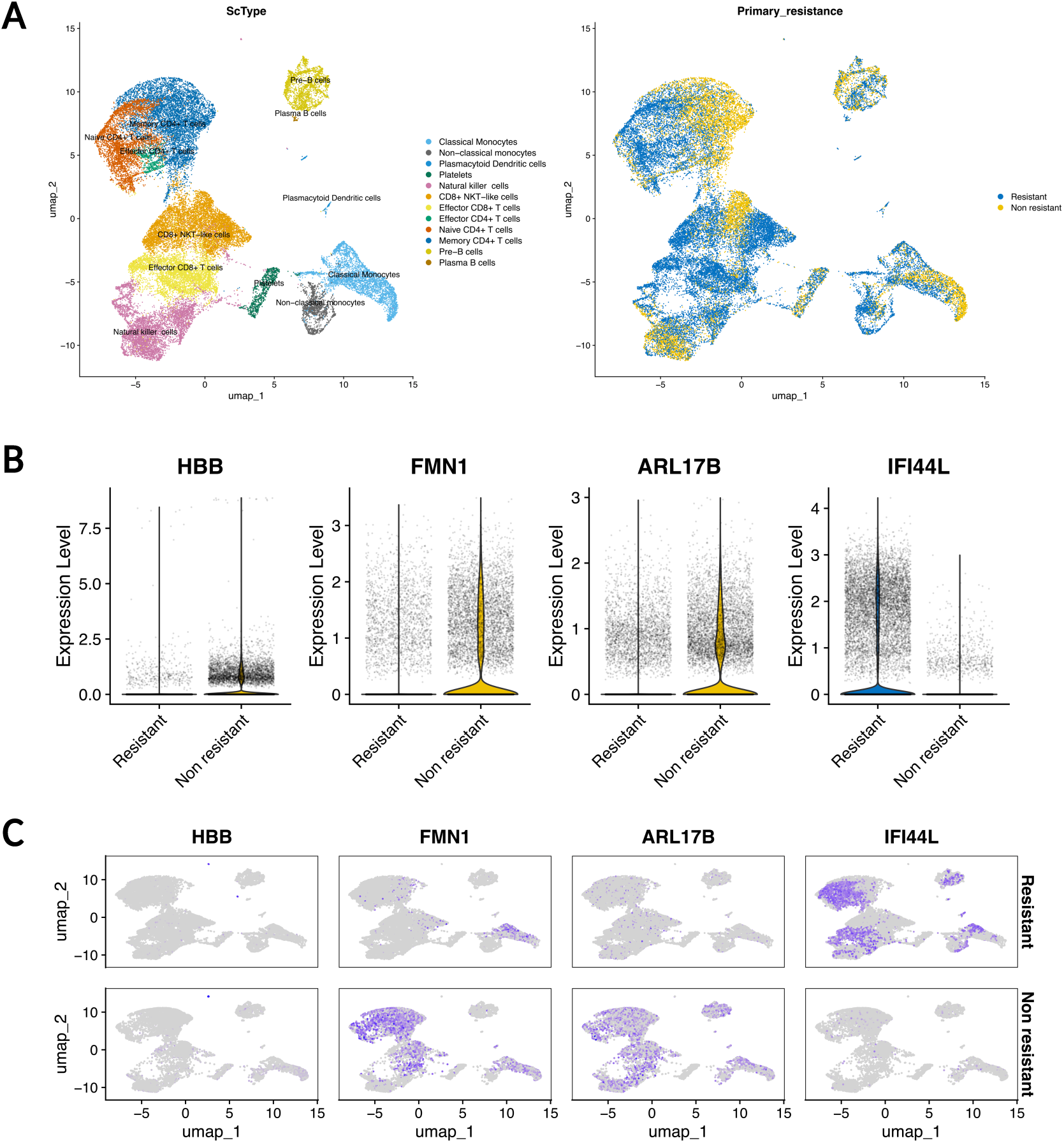
Single-cell analysis and DE genes linked to ICI primary resistance. (A) UMAP visualization of scRNA-seq data showing 12 distinct immune cell types inferred using sc-type. (B) UMAP with the primary resistance annotation. (C) Violin plot displaying the expression levels of the 4 DE genes identified when comparing resistant patients and non-resistant patient in the Single-cell sub-cohort. (D) UMAP plot with the colour based on the expression of the 4 DE genes.

After preprocessing, normalization, and cell type annotation, a differential expression analysis with the scRNA-seq data was performed based on primary resistance. This analysis included 3 non-resistant patients and 5 resistant patients (figure 4B). As a result, only 4 DE genes were identified, with 3 upregulated in non-resistant patients (*HBB, FMN1, ARL17B*) and 1 upregulated in resistant patients (*IFI44L*) (figure 4C). Except for *HBB*, the remaining genes exhibit a relatively homogeneous distribution across the entire UMAP of the group of patients expressing the gene (figure 4D).

Leveraging the resolution of single-cell sequencing and cell identity annotation, we next assessed primary resistance at the individual cell type level. Differential expression analysis was conducted for each cell type by comparing cells from resistant and non-resistant patients, leading to the identification of a set of DE primary resistance genes specific to each cell type (supplementary table S3). Classical Monocytes, Naive CD4^+^ T cells, Non-classical Monocytes, and CD8^+^ NKT-like cells exhibited the highest number of DE genes when comparing resistant and non-resistant patients, with 40, 35, 15, and 13 DE genes, respectively (figure 5A).

**Figure 5.**
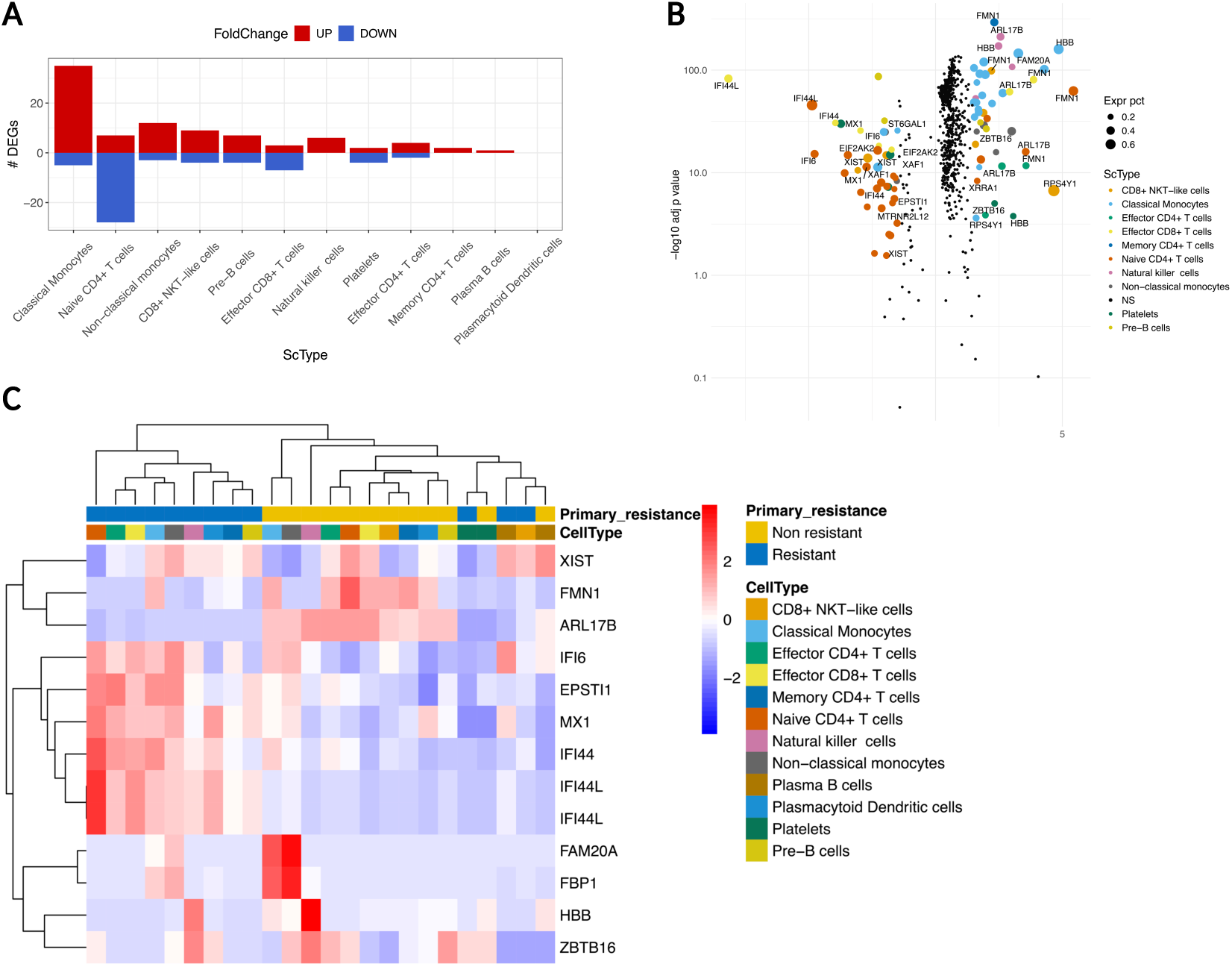
DE analysis of primary resistance across immune cell types. (A) Number of DE genes associated with primary resistance across each annotated cell type in the single-cell analysis. (B) Volcano plot highlighting genes that are present in more than one cell type, with point size representing the percentage of cells expressing each gene. (C) Heatmap displaying the average gene expression differences across cell types stratified by primary resistance and cell type, with hierarchical clustering applied to both rows and columns. NS: not significant.

Across all individual analyses, we observed a subset of genes that exhibit significant differences in response across different cell types (figure 5B). *ARL17B, FAM20A, FBP1, FMN1, HBB*, and *ZBTB16* emerged as candidate pan cell-type genes associated with non-resistance (supplementary figure S3), whereas *IFI44, IFI44L, IFI6, EPSTI1*, and *MX1* were linked to resistance across multiple cell types (supplementary figure S4). *XIST* and *RPS4Y1*, both encoded on sex chromosomes, were found to be significantly associated with resistant and non-resistant status respectively, despite the inclusion of sex as a covariate in the analysis. Their persistent signal suggests a potential biological role independent of sex bias. Further validation in a larger cohort stratified by sex will be essential to confirm their relevance as biomarkers.

Therefore, by performing differential expression analysis per and across cell subtypes, we expanded the blood gene signature associated with primary resistance from 4 (*HBB, FMN1, ARL17B and IFI44L*) to 13 genes (*HBB, FMN1, ARL17B, IFI44L, FAM20A, FBP1, ZBTB16, IFI44, IFI6, EPSTI1, MX1* and XIST) in the single-cell analysis (figure 5C).

When the stratification in non-resistant and resistant patients was attempted using the cell type abundance in this Single-cell sub-cohort, we did not find any significant associations, likely due to the low number of patients in this analysis (3 non-resistant vs. 5 resistant). However, the cell identity markers identified in the beginning of this analysis were integrated as a reference to perform deconvolution on the bulk RNA-seq tissue samples (supplementary table S4) in the multidimensional analysis.

### Multidimensional analysis to reveal cellular identity markers of primary resistance

Single-cell RNA sequencing enabled us to obtain the transcriptomic profiles of each detected immune blood circulating cell type. Using this scRNA-seq data, we identified cell-type specific markers (supplementary table S4) and constructed a deconvolution matrix. This matrix was then applied to the solid tumor bulk RNA-seq samples from the Discovery FFPE bulk-RNA seq sub-cohort, allowing us to accurately estimate the relative abundance of each immune cell population within the TIME (figure 6A) (supplementary table S5). Furthermore, the same workflow was applied to the Validation cohort (supplementary table S6). The relative abundance estimated through deconvolution analysis in solid tumor samples was compared between non-resistant and resistant patients within the Discovery FFPE bulk-RNA seq sub-cohort (figure 6B) and the Validation cohort (figure 6C). Higher proportions of Plasma cells, Pre-B cells, Memory CD4^+^ T cells, and Naive CD4^+^ T cells were observed in the TIME of non-resistant patients compared to resistant patients.

**Figure 6.**
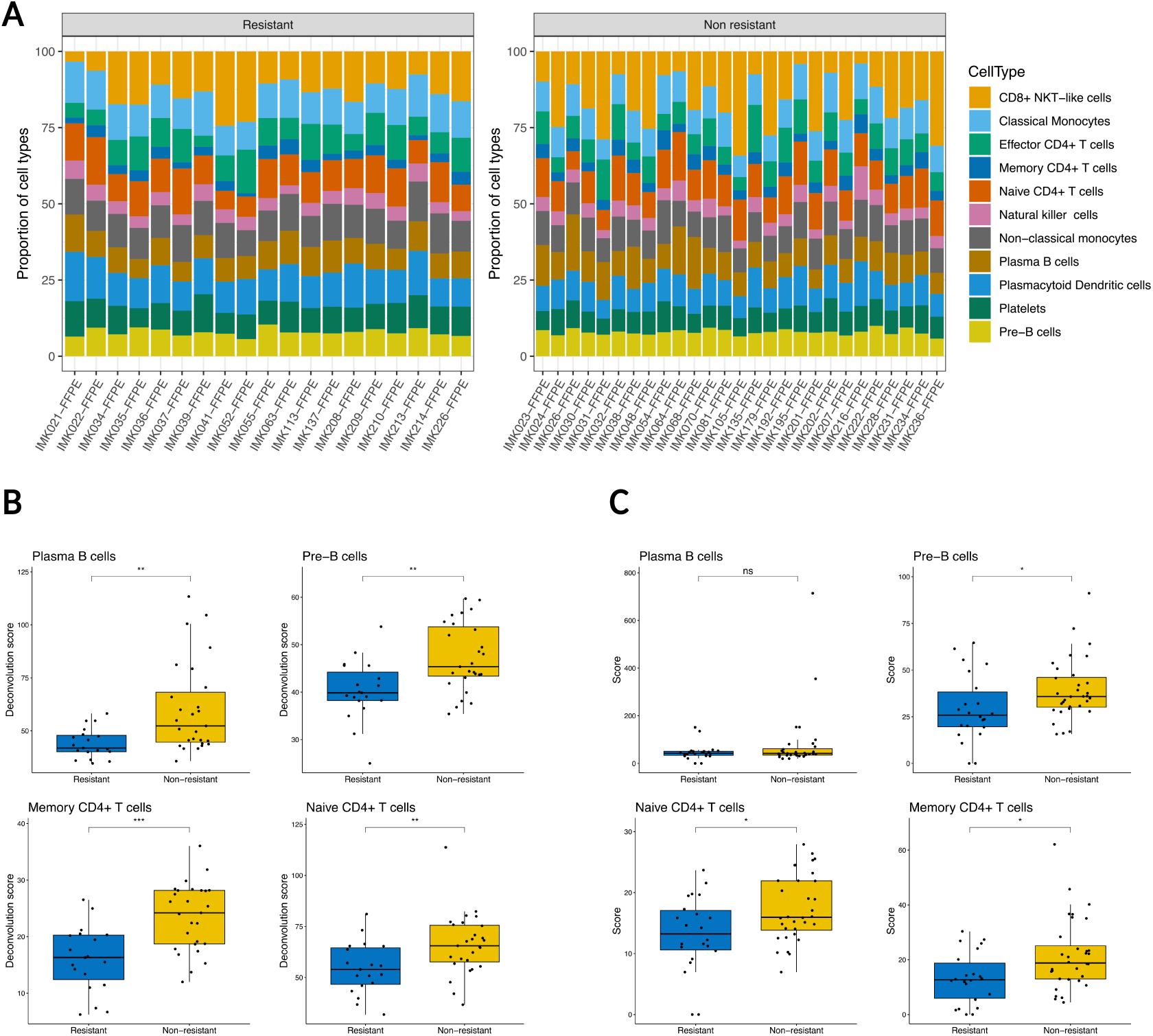
Bulk RNA-Seq deconvolution analysis of immune cell types in primary resistance. (A) Bar plot showing the percentage of inferred immune cells from bulk RNA-seq data in the Discovery FFPE bulk-RNA seq sub-cohort, stratified by primary resistance status, based on the cell types annotated in the single-cell analysis. (B) Deconvolution scores of significant cell type abundance differences derived from bulk RNA-seq data between non-resistant and resistant patients in the Discovery FFPE bulk-RNA seq sub-cohort. (C) Deconvolution analysis applied to the Validation cohort. Statistical significance is indicated as: ns (non-significant), * (p < 0.05), ** (p < 0.01), *** (p < 0.001).

The predominant markers in these cell populations are *TSHZ2* and *TRABD2A* in Naive CD4^+^ T cells, whereas *TESPA1* and *TOMM7* in Memory CD4^+^ T cells, and *IL7R* as a shared marker in both CD4^+^ T cell types. In Plasma cells, the main markers are *XBP1* and *MZB1*, while in Pre-B cells, *BLK*, *MS4A1, EBF1* and *PAX5* are the key markers (supplementary table S4, supplementary figure S5, supplementary table S8).

### Prognosis signature based on an immune score identifies a cohort of long-term survivors

The estimation of immune cell populations within the TIME identified key components implicated in the response to ICIs, including Naive and Memory CD4^+^ T cells, Pre-B cells, and Plasma cells. Given their association with primary resistance, we next assessed their prognostic value in determining long-term patient overall survival. This is possible given the long follow-up of the patients included in the different cohorts, with median follow-up times of 32.55 months (95% CI: 29.61, 55.05) in the Discovery FFPE bulk-RNA seq sub-cohort, 28.13 months (95% CI: 26.57, 30.57) in the Extended Validation cohort, and 32.29 months (95% CI: 24.98, 49.80) in the Discovery Flow cytometry sub-cohort. When combined into a single composite score, these immune components emerge as a promising prognostic signature. As shown in figure 7, the ImmuneSignature, derived from the estimation of the four cell types (Plasma cells, Pre-B cells, Memory CD4^+^ T cells, and Naive CD4^+^ T cells) in the tumor microenvironment of the Discovery FFPE bulk-RNA seq sub-cohort (figure 7A), significantly separates the survival curves. Patients with high abundance of these immune components show a survival indicative of a better prognosis (survival median 34 months (27 months - NR)), while those with lower abundance experience poorer outcomes (3 months, (1.77 months - NR)). When validated in the Extended Validation cohort of 268 patients (figure 7B), this prognostic signature exhibits a consistent pattern, with better prognosis in those with a high ImmuneSignature (NR), and worse prognosis in those with a lower ImmuneSignature (21 months, (14 months-27 months)).

**Figure 7.**
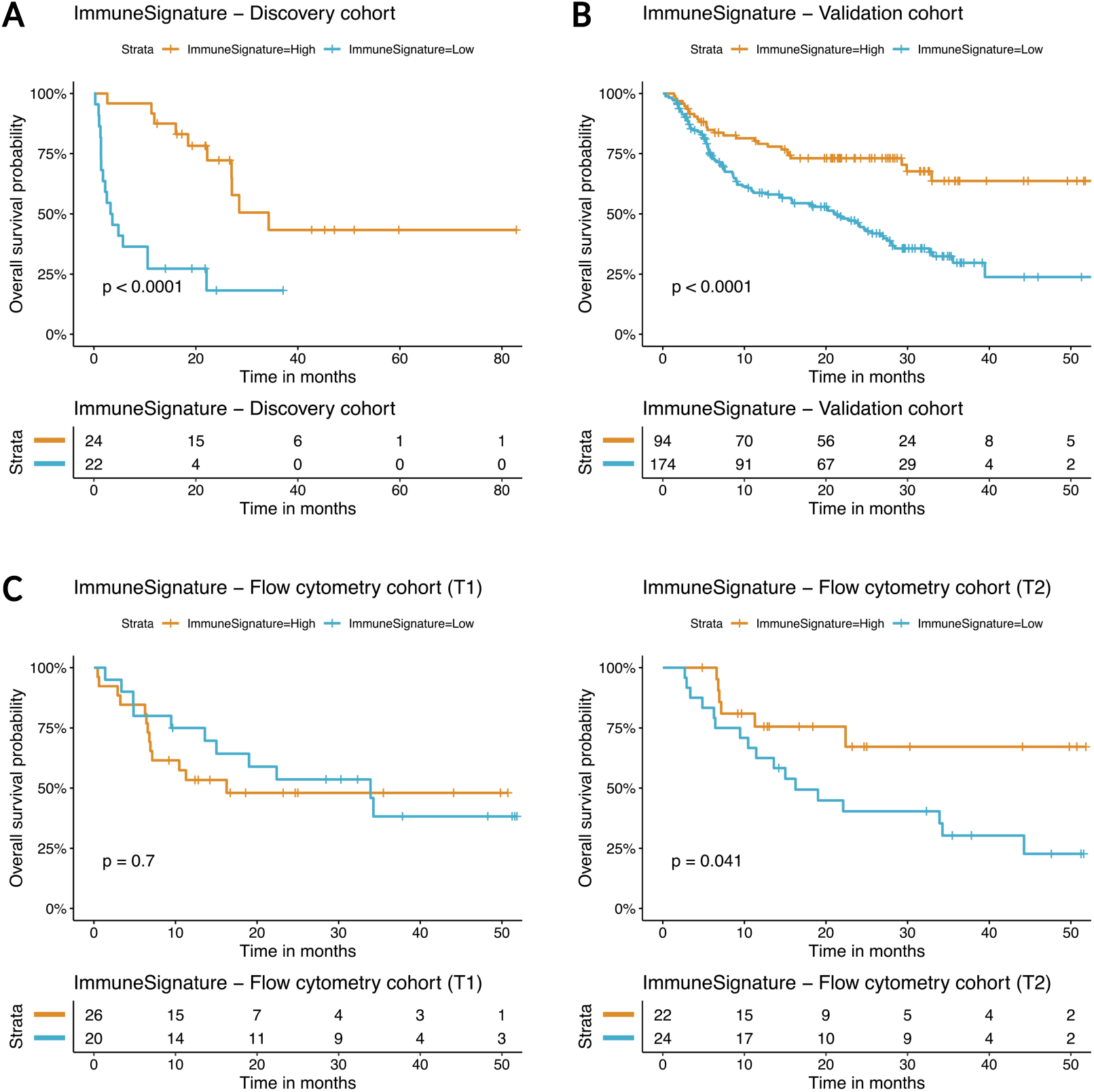
Overall survival analysis stratified by ImmuneSignature. Kaplan-Meier survival curves illustrating the impact of immune signatures derived from B cell and CD4^+^ T cell deconvolution analysis on overall survival. (A) Survival analysis in the Discovery FFPE bulk-RNA seq sub-cohort, showing significant stratification based on ImmuneSignature levels. (B) Validation of the ImmuneSignature’s prognostic value in the independent Extended Validation cohort. (C) Kaplan-Meier survival curves from the Flow cytometry cohort, based on the ImmuneSignature assessed at T1 (prior to the initiation of immunotherapy) and T2 (after the second treatment cycle). Statistical significance was assessed using the log-rank test.

Additionally, to evaluate this score as a surrogate in liquid biopsy, we analyzed CD4^+^ T cell, comprising Naive and Memory CD4^+^ T cells, and Plasma B cell abundance by flow cytometry. As only a subset of the four relevant immune cell types were available for phenotyping by flow cytometry, this analysis represents a partial reconstruction of the proposed ImmuneSignature. We calculated the score for T1 (prior to the initiation of treatment) and T2 (after the second cycle) in the Discovery Flow cytometry sub-cohort (n=46) (figure 7C). Patients were categorized into high ImmuneSignature and low ImmuneSignature groups independently based on markers values at T1 and T2. At T1, no significantly separate survival curves were observed. However, at T2, patients with high ImmuneSignature had a median OS of 22.40 months (NR-NR), while those with low ImmuneSignature had a median OS of 16.26 months (11-NR). In T2, the Log-rank test comparing the survival curves yielded a p-value < 0.05, indicating a statistically significant difference between the two groups.

## Discussion

The identification of patients likely to exhibit primary resistance remains a critical challenge for the optimized use of ICIs. To date, transcriptomic analyses have not identified a predictive or prognostic signature for primary resistance (4) in an homogenous cohort of metastatic cutaneous melanoma involving tissue and liquid biopsy samples. In this study, we compared patients with primary resistance to anti-PD-1. Our findings, validated in external cohorts, generate promising molecular and cellular markers that address this gap.

The analysis of tumor samples from a cohort of 46 patients identified an 82-gene transcriptomic signature of coding and non-coding transcripts. Several of these genes have been previously associated with anti-tumoral activity or tumor progression. *GRIK3* has been proved as a marker of poor prognosis in diverse cancers due to its role in oncogenic processes such as proliferation, cellular migration, and epithelial-to-mesenchymal transition (EMT) (20). Its overexpression in melanoma patients exhibiting primary resistance to ICIs aligns with its prognostic significance in other malignancies, as previously reported (21). Interestingly, we also identified the long non-coding RNA *LINC01502*, which has been linked to BRAF inhibitor treatment resistance in melanoma (22). Regulatory networks involving lncRNA–miRNA–mRNA interactions reported for this transcript implicate the transcription factor MITF, a central regulator in melanoma biology. Patients with elevated levels of *LINC01502* display the more aggressive phenotype, potentially due to its function as a competitive regulator of miRNAs shared with MITF (23). Our findings indicate that *LINC01502* is a key modulator of the tumor microenvironment, contributing to disease progression and limiting the efficacy of ICIs. Notably, its association with progression observed in our study is consequent with its previously reported role in the literature as a biomarker of resistance to BRAF inhibitors, further supporting its relevance across different therapeutic contexts (22).

Moreover, we identified genes upregulated in non-resistant patients already reported as prognostic biomarkers for immunotherapy outcomes in other cancers, such as *FDCSP* in kidney renal clear cell carcinoma (24). Furthermore, out of the 60 upregulated genes, 36 were previously reported in a previous cohort by JL Onieva et al. (2022) (25), with overlapping genes primarily linked to the B-cell lineage. Notably, these include immunoglobulin-related genes such as *IGKC* and the *IGHG* family. The involvement of these immunoglobulin-related genes underscores their potential role in the humoral immune response, particularly in processes such as clonal expansion and affinity maturation (26,27). The upregulated gene *WDR63* emerged as the most significant variable of primary resistance in our predictive model, constructed from the differential expression of genes from bulk RNA-seq data. *WDR63* contributes to metastasis suppression, acting as a transcriptional target of p53 (28). Functionally, *WDR63* mediates p53’s anti-metastatic effects by inhibiting *Arp2/3-mediated actin polymerization*, a process essential for cell migration and invasion (29). This suggests that *WDR63* plays a pivotal role in preventing tumor progression and promoting a more favorable therapeutic outcome.

In addition to genes associated with the B-cell lineage, our signature also includes genes that functionally link B and T cells, such as *CXCL13* (30,31). The CXCL13/CXCR5 axis is central in orchestrating immune processes mediated by B and T cells, particularly in the context of antitumor defense. *CXCL13* is a chemokine induced in exhausted CD8^+^ T cells under the influence of TGF-β. It promotes the recruitment of follicular B-helper CD4^+^ T cells and B cells to the tumor microenvironment thereby fostering inflammation, enhancing the accumulation of other immune cell populations, and facilitating the formation of tertiary lymphoid structures (TLS) (32). The presence of *CXCL13* in our transcriptomic signature emphasizes the importance of B-T cell crosstalk in shaping a robust anti-tumor immune response and underscores its potential as a biomarker for favorable ICIs outcomes (33), as also supported by findings from spatial transcriptomics studies (34).

To elucidate the biological function underlying resistance, we conducted a pathway analysis of genes overexpressed in resistant patients. Notably, two critical signaling pathways: *IL-12 signaling and production in Macrophages* and *CREB signaling*, were significantly activated in primary resistant cases. The inhibition of IL-12 signaling in non-resistant cases appears to correlate with the activation of immune pathways centered on B cell-mediated responses. As discussed before, B cell functions might contribute significantly to overcoming tumor progression through an integrated innate-adaptive immune response, aimed at controlling metastasis. On the other hand, the activation of *CREB signaling* in primary resistant melanoma cases points to a potential mechanism of immune evasion and tumor persistence. CREB, a key transcription factor involved in the regulation of immune responses and inflammation, has been shown to promote pathways that can dampen the immune system’s effectiveness (35). The activation of CREB could, therefore, facilitate immune suppression and reduce the therapeutic success of immunotherapy.

When examining the expression of specific genes in single-cell RNA-seq data, several genes were associated with non-resistance to immunotherapy. *FMN1* has been linked as a key predictor of PD-1 inhibitor response in glioblastoma, reflecting an immune-activating tumor microenvironment (36). In our analysis, *FMN1*, was upregulated in the non-resistant Memory CD4^+^ T cells, Naive CD4^+^ T cells, Effector CD4^+^ T cells, Effector CD8^+^ T cells and CD8^+^ NKT-like cells. Also, our data shows that melanoma patients who responded to immunotherapy exhibited significant overexpression of *ZBTB16* in both Effector CD4^+^ T cell and Pre-B cells. Given the key regulatory function of ZBTB proteins in T cell development, differentiation, and effector function (37), *ZBTB16* could enhance the functional capacity of these immune cells. This upregulation may promote a more robust and coordinated anti-tumor response, whereas its downregulation in non-responders could contribute to diminished immune activation and primary resistance to immunotherapy.

The genes *IFI44, IFI44L, IFI6*, and *MX1* were identified as markers associated with resistance in the single-cell analysis. *IFI44* and *IFI44L* have been associated with aggressive melanoma subtypes characterized by invasive, de-differentiated gene expression patterns (38). Furthermore, *IFI6* – in squamous cell carcinoma (39) – and *MX1* – in a mouse model (40) – have been shown to promote immune and facilitate tumor progression, indicating their role in immune escape mechanisms.

Finally, we explored the expression of *XIST* and *RPS4Y1*, two genes located on sex chromosomes, in the context of immunotherapy response. Although sex was included as a covariate in our analytical model, both genes remained significantly associated with clinical outcome, suggesting a potential biological role independent of sex-related confounding. These findings warrant validation in larger cohorts with sufficient power for sex-stratified analyses. *XIST* was overexpressed in non-responders, particularly within CD8^+^ NKT-like cells and Classical Monocytes. This observation aligns with prior studies indicating that *XIST* functions as a context-dependent immunoregulatory element, modulating X chromosome inactivation and gene silencing in lymphocytes (41). Notably, its expression is dynamically regulated during T cell activation, and persistent upregulation in specific immune subsets may reflect an exhausted or immunosuppressive phenotype associated with ICI resistance. Conversely, *RPS4Y1* was found to be overexpressed in responders within the same immune compartments. This gene has previously been linked to enhanced immune infiltration and favorable responses to immune checkpoint blockade in non-small cell lung cancer (42).

Given the biological relevance of the cellular distribution within the tumor microenvironment, we performed a deconvolution analysis of the FFPE RNA-seq data using specific markers of various cell populations circulating in patients with the same histology that were to be treated with ICIs (15). This analysis revealed an underrepresentation of Pre-B cells, Plasma cells, Naive CD4^+^ T cells, and Memory CD4^+^ T cells in patients with primary resistance.

The role of CD4^+^ T cells in antitumor responses has been extensively reviewed (43,44). CD4^+^ T cells, essential in orchestrating effective antitumor immunity, engage in complex interactions with various cells within the tumor microenvironment, including myeloid-derived suppressor cells (43). Notably, Memory CD4^+^ T cells have been implicated in improved survival outcomes with ICIs. The importance of these cells in ICIs-based immunotherapy lies in their resistance to activation-induced cell death and their ability to differentiate into an effector memory phenotype, significantly contributing to clinical responses (44).

In addition to our findings regarding CD4^+^ T cells, our study again sheds light on the relevance of B cells in the context of immune checkpoint inhibition in metastatic cutaneous melanoma. The contribution of B cell–mediated humoral immunity is reinforced through our integrative multidimensional analysis. In the bulk RNA-seq data, BCR signaling was the most significantly enriched pathway in non-resistant patients, aligning with the observed expansion of Pre-B and Plasma cell populations. Notably, *MZB1* emerged as a central integrative marker across both bulk and single-cell transcriptomic dimensions. It was identified as a key variable in our predictive model and also serves as a specific marker for both B cell subsets, which are enriched in our ImmuneSignature. In support of our findings, *MZB1* has been described as a B cell–specific endoplasmic reticulum chaperone involved in antibody secretion and immunoglobulin folding. Its expression is closely linked to Plasma cell function and B cell activation (45) and it has also been shown to correlate with B cell clonal expansion (46), reinforcing its role in active humoral responses. Additionally, it has been reported to influence the immune microenvironment and reduce ovarian cancer cell migration (47), highlighting its potential function at the interface between B cell–mediated immunity and tumor progression. Interestingly, Plasma cells are one of the PBMCs subpopulations that have validated the ImmuneSignature by flow cytometry (supplementary table S8).

While previous studies have demonstrated the prognostic and mechanistic significance of intratumoral B cells in melanoma immunotherapy response (25,48,49) none have integrated both tissue-level and peripheral blood data within a histologically and therapeutically homogeneous cohort of patients treated with the anti–PD-1 axis and classified according to SITC-defined primary resistance criteria (5). Of particular relevance in the context of ICIs, a recent large-scale study in head and neck squamous-cell carcinoma further reinforces the importance of the abundance of both intratumoral and circulating B cells as a strong predictor of ICI response underscoring their potential utility as accessible non-invasive biomarkers (50). In comparison, that study included a heterogeneous cohort encompassing multiple disease subtypes and treatment regimens, whereas our study focuses on a clinically homogeneous population enabling a more controlled and specific evaluation of immune biomarkers associated with primary resistance. Overall, these aspects represent key strengths of our study and highlight its added value within the evolving landscape of immuno-oncology research. However, additional mechanistic and experimental studies will be required to fully elucidate the roles of B cells in antitumor immunity particularly in relation to antibody secretion, antigen presentation and cytokine-mediated modulation.

Our study evaluates the prognostic and predictive capabilities of the findings presented. To date, several transcriptomic studies have explored biomarkers of response to ICIs in melanoma; however, significant limitations persist across these external datasets. Among the seven publications with analysis of melanoma patients treated with ICI (7,9–14), including the Validation cohort (7) and the six cohorts conforming the Extended Validation Cohort (7,9–13), a recurring limitation is the inclusion of heterogeneous melanoma subtypes—most commonly a mixture of cutaneous, acral, and mucosal melanomas, which exhibit distinct biological profiles. Moreover, external validation is generally limited or absent in these analyses. For example, Gide et al. (2019) (9) relied exclusively on training performance without applying any external validation. Liu et al. (2019) (7) reported validation on a minimal sample size (n = 8), using 10-fold cross-validation rather than an independent cohort. The remaining studies exhibit further limitations: Hugo et al. (2016) (10) and Riaz et al. (2017) (12), did not report a predictive model or provide standard performance metrics, while Li et al. (2023) (14) based their evaluation solely on OS, without assessing predictive discrimination. Only Cui et al. (2021) (11) included a large and independent validation cohort. Despite these limitations, important biological insights have emerged from them: higher mutational burden (Hugo et al. (2016) (10)), neoantigen load, and cytolytic activity (Van Allen et al (2015) (13)) have been associated with improved clinical outcomes; other findings include enrichment of mesenchymal transition and extracellular matrix (Hugo et al. (2016) (10)) remodeling pathways in non-responders, and the presence of specific immune cell phenotypes such as EOMES^+^CD69^+^ effector memory T cells in responders (Gide et al. (2019) (9)), B cell enrichment (Liu et al. (2019) (14)) and T cell-inflamed tumor microenvironment (Cui et al. (2021) (11)). In contrast, our study leverages a homogeneous cohort of patients with metastatic cutaneous melanoma, systematically evaluating both primary resistance and OS in the same population. By focusing exclusively on cutaneous melanoma, we reduce biological variability and enhance the relevance and specificity of transcriptional predictors.

Other studies have mainly developed predictive models using tissue (51) or liquid biopsy data (52). TIDE algorithm with n=25, AUC of 0.83 and 770 genes (51) and anti-PD1 liquid biopsy cohort in (52) n=40, AUC of 0.57 and only one cell type candidate. To address the challenge of limited generalizability, our predictive signature was independently validated in a cohort of 54 metastatic cutaneous melanoma patients, demonstrating robust performance with an AUC of 0.841. This approach underscores the relevance role of comprehensive molecular profiling in improving clinical outcomes, as demonstrated in other precision oncology frameworks, Kerle et al. (2025) (53). Moreover, beyond its predictive capacity, the identification of specific cellular populations has highlighted a subset of patients with long-term overall survival that generates a new hypothesis as a non-invasive surrogate score. Further validation of these findings could lead to the development of a marker-based panel, enhancing the ability to refine patient stratification and inform therapeutic decisions.

Although our findings support the predictive potential of multi-gene immune signatures, we acknowledge the practical challenges associated with their implementation in routine clinical settings. These include assay complexity, cost, turnaround time, and the need for standardization across platforms. While our study did not directly address spatial dynamics, these findings raise important questions regarding the localization and functional role of the highlighted immune cells within the tumor microenvironment. Future studies incorporating spatial transcriptomics or imaging-based techniques will be essential to elucidate these spatial relationships and their relevance to immunotherapy resistance. Regarding the surrogate ImmuneSignature assessed in liquid biopsy, our study did not employ customized flow cytometry panels specifically designed to capture the four immune cell populations comprising the original signature. However, the use of available markers enabled meaningful proof-of-concept validation. Notably, this approach successfully stratified patients by survival in samples collected at T2, 2 or 3 weeks after ICI initiation, supporting the translational relevance of the identified subrogate ImmuneSignature as an early on-treatment biomarker. Further refinement of this technique, particularly using flow cytometry, a method already established in clinical practice, could represent a significant step forward in translating immune-based signatures into actionable tools for patient management. Collectively, our approach not only serves as a model for patient selection but also provides insights into a potential molecular profiling framework to further explore therapeutic targets. This could facilitate the development of novel combination strategies with ICIs, as previously proposed in the literature by incorporating adoptive T-cell therapy (54).

## Conclusions

In conclusion, this study identifies and validates a robust transcriptomic signature as a promising tool for predicting primary resistance to ICIs in metastatic cutaneous melanoma. Our findings underscore the pivotal role of B cell–mediated mechanisms and the complex interplay between innate and adaptive immune responses in shaping resistance to immunotherapy. Through integrative, multidimensional analyses, we uncovered key resistance-associated genes (*CXCL13, WDR63, MZB1, FDCSP, IGKC* and *GRIK3*), and circulating immune cell types (Plasma cells, Pre-B cells, Memory CD4^+^ T cells, and Naive CD4^+^ T cells) that together define an ImmuneSignature measurable via both bulk RNA-seq and flow cytometry. Together, these insights pave the way for the development of resistance-targeted therapeutic strategies and offer a precision oncology framework to optimize treatment selection in cutaneous melanoma.

## Study limitations

Several limitations should be acknowledged in this study. First, the predictive model developed using tissue samples would benefit from validation in a larger cohort, which could strengthen its generalizability, as it has been possible for the ImmuneSignature prognosis score. Additionally, the inclusion of paired molecular data would enrich the study, providing further insights into the relationship between different molecular profiles. Although single-cell RNA sequencing was performed with high cellular depth, the sample size was limited to eight patients. While this provided valuable resolution at the cellular level, a larger cohort will be essential to validate and extend the observed findings. In this context, the deconvolution of bulk RNA-seq data provides a transcriptional snapshot of circulating immune cells but does not fully account for the complexity of the tumor immune microenvironment, including spatial organization and tissue-resident immune populations that may critically influence therapeutic response. Lastly, although flow cytometry was utilized to validate the ImmuneSignature, only an approximate representation of the four populations determining tissue-based ImmuneSignature were distinguishable with the available data in our cohort, which may affect the precision of cellular profiling. This limitation reduces the resolution of the validation and may affect the ability to capture the full complexity of the signature. A more comprehensive panel including additional lineage and activation markers, or the use of higher-dimensional technologies, will likely be required to validate the signature.

## Abbreviations

AUC: Area Under the Curve
BCR: B Cell Receptor
DE: Differential Expression
EMT: Epithelial-Mesenchymal Transition
FFPE: Formalin-Fixed Paraffin-Embedded
GEMs: Gene Expression Modules
GEO: Gene Expression Omnibus
ICIs: Immune Checkpoint Inhibitors
MSI: Microsatellite Instability
NR: Not reached
OS: Overall Survival
PBMCs: Peripheral Blood Mononuclear Cells
PFS: Progression-Free Survival
RNA-seq: RNA Sequencing
ROC: Receiver Operating Characteristic
scRNA-seq: Single-Cell RNA Sequencing
TLS: Tumor Lymphoid Structures
TIME: Tumor Immune Microenvironment
TMB: Tumor Mutational Burden
UMAP: Uniform Manifold Approximation and Projection

## Declarations

### Ethics approval and consent to participate

The study follows the Declaration of Helsinki and has been submitted and approved by Comité de Ética de la Investigación Provincial de Málaga. Approval date on 26 October 2017 with the title: “Omics integration for precision cancer immunotherapy” (799818, H2020-MSCA-IF-2017) research project. All patients signed Informed Consent to participate in the study and received an information sheet about the project.

### Consent for publication

Not applicable.

## Data availability

The original contributions presented in the study are included in the article Supplementary Material and Supplementary Table, further inquiries can be directed to the corresponding author. Data is available upon reasonable request. All data analyzed in our study are available upon request.

## Code availability

Code available upon request on http://github.com/ImmunoOncology.

## Competing interests

The authors declare no conflict of interest.

## Funding

The work is funded by Instituto de Salud Carlos III through the projects PI18/01592 (Co-funded by the European Regional Development Fund/European Social Fund “A way to make Europe”/“Investing in your future”), PI22/01816 (Co-funded by the European Union), and DTS23/00114, Sociedad Española de Oncología Médica (SEOM); Sistema Andaluz de Salud, through the projects SA 0263/2017 and RC-0009-2021 (Nicolás Monardes); Consejería de Transformación económica, Industria, Conocimiento y Universidades through the projects PREDOC-01691 and ProyExcel_01002; Spanish Group of Melanoma (Award for Best Research Project 2023), Agencia Estatal de Investigación, Consolidación Investigadora (CNS2023-145629) and Asociación Española Contra el Cáncer, Proyectos Estratégicos (PRYES247250BARR).

## Author contributions

**J.L. O.**: Conceptualization, investigation, formal analysis, data curation, and writing – original draft. **E. P.-R.**: Conceptualization, resources, and supervision. **V. V.**, **L. F.-O.**, and **B. M.-G.**: Formal analysis. **M. B.-G**. and **M. Z.**: Resources. **I. B.**: Conceptualization, investigation, supervision, writing – review and editing, and funding acquisition. **A. R.-D.**: Investigation, supervision, and writing – review and editing. All authors reviewed and approved the final manuscript.

## Acknowledgments

The authors gratefully acknowledge the Supercomputing and Bioinnovation Center (SCBI) at the University of Malaga for providing computational resources (http://www.scbi.uma.es/). They are also grateful to the Hospital-IBIMA Biobank (Biobanco del Sistema Sanitario Público de Andalucía) integrated in the Spanish National biobanks Network (PT23/00049) supported by ISCIII. Additionally, the authors express their gratitude to Andrea González, Marina Rivero, Alberto Ríos, Alba Rodríguez and Javier Oliver for their valuable support and insightful suggestions, which significantly enhanced the quality of this work.

## Notes

### Competing Interest Statement

The authors have declared no competing interest.

## References

1. Robert C, Long GV, Brady B, Dutriaux C, Maio M, Mortier L, et al. Nivolumab in previously untreated melanoma without BRAF mutation. N Engl J Med. 2015 Jan 22;372(4):320–30.

2. Mok TSK, Wu YL, Kudaba I, Kowalski DM, Cho BC, Turna HZ, et al. Pembrolizumab versus chemotherapy for previously untreated, PD-L1-expressing, locally advanced or metastatic non-small-cell lung cancer (KEYNOTE-042): a randomised, open-label, controlled, phase 3 trial. Lancet Lond Engl. 2019 May 4;393(10183):1819–30.

3. Sharma P, Allison JP. Immune checkpoint targeting in cancer therapy: toward combination strategies with curative potential. Cell. 2015 Apr 9;161(2):205–14.

4. Sun Q, Hong Z, Zhang C, Wang L, Han Z, Ma D. Immune checkpoint therapy for solid tumours: clinical dilemmas and future trends. Signal Transduct Target Ther. 2023 Aug 28;8(1):320.

5. Kluger HM, Tawbi HA, Ascierto ML, Bowden M, Callahan MK, Cha E, et al. Defining tumor resistance to PD-1 pathway blockade: recommendations from the first meeting of the SITC Immunotherapy Resistance Taskforce. J Immunother Cancer. 2020 Mar 1;8(1):e000398.

6. Lei Y, Li X, Huang Q, Zheng X, Liu M. Progress and Challenges of Predictive Biomarkers for Immune Checkpoint Blockade. Front Oncol. 2021 Mar 11;11:617335.

7. Liu D, Schilling B, Liu D, Sucker A, Livingstone E, Jerby-Arnon L, et al. Integrative molecular and clinical modeling of clinical outcomes to PD1 blockade in patients with metastatic melanoma. Nat Med. 2019 Dec;25(12):1916–27.

8. Alvarez-Breckenridge C, Markson SC, Stocking JH, Nayyar N, Lastrapes M, Strickland MR, et al. Microenvironmental Landscape of Human Melanoma Brain Metastases in Response to Immune Checkpoint Inhibition. Cancer Immunol Res. 2022 Aug 3;10(8):996–1012.

9. Gide TN, Quek C, Menzies AM, Tasker AT, Shang P, Holst J, et al. Distinct Immune Cell Populations Define Response to Anti-PD-1 Monotherapy and Anti-PD-1/Anti-CTLA-4 Combined Therapy. Cancer Cell. 2019 Feb 11;35(2):238–255.e6.

10. Hugo W, Zaretsky JM, Sun L, Song C, Moreno BH, Hu-Lieskovan S, et al. Genomic and Transcriptomic Features of Response to Anti-PD-1 Therapy in Metastatic Melanoma. Cell. 2016 Mar 24;165(1):35–44.

11. Cui C, Xu C, Yang W, Chi Z, Sheng X, Si L, et al. Ratio of the interferon-γ signature to the immunosuppression signature predicts anti-PD-1 therapy response in melanoma. NPJ Genomic Med. 2021 Feb 4;6(1):7.

12. Riaz N, Havel JJ, Makarov V, Desrichard A, Urba WJ, Sims JS, et al. Tumor and Microenvironment Evolution during Immunotherapy with Nivolumab. Cell. 2017 Nov 2;171(4):934–949.e16.

13. Van Allen EM, Miao D, Schilling B, Shukla SA, Blank C, Zimmer L, et al. Genomic correlates of response to CTLA-4 blockade in metastatic melanoma. Science. 2015 Oct 9;350(6257):207–11.

14. Li A, Luo L, Du W, Yu Z, He L, Fu S, et al. Deciphering transcriptomic determinants of the divergent link between PD-L1 and immunotherapy efficacy. Npj Precis Oncol. 2023 Sep 11;7(1):1–13.

15. Newman AM, Steen CB, Liu CL, Gentles AJ, Chaudhuri AA, Scherer F, et al. Determining cell-type abundance and expression from bulk tissues with digital cytometry. Nat Biotechnol. 2019 Jul;37(7):773–82.

16. Chiu YJ, Ni CE, Huang YH. HArmonized single-cell RNA-seq Cell type Assisted Deconvolution (HASCAD). BMC Med Genomics. 2023 Oct 31;16(2):272.

17. Hao Y, Stuart T, Kowalski MH, Choudhary S, Hoffman P, Hartman A, et al. Dictionary learning for integrative, multimodal and scalable single-cell analysis. Nat Biotechnol. 2024 Feb;42(2):293–304.

18. Nader K, Tasci M, Ianevski A, Erickson A, Verschuren EW, Aittokallio T, et al. ScType enables fast and accurate cell type identification from spatial transcriptomics data. Bioinforma Oxf Engl. 2024 Jul 1;40(7):btae426.

19. Onieva JL, Cháves P, Oliver J, Garrido-Barros M, Zafra J, Sojo B, et al. Abstract 1910: flowTOTAL: A comprehensive bioinformatics workflow for flow cytometry automatic analysis. Cancer Res. 2022 Jun 15;82(12_Supplement):1910.

20. Xiao B, Kuang Z, Zhang W, Hang J, Chen L, Lei T, et al. Glutamate Ionotropic Receptor Kainate Type Subunit 3 (GRIK3) promotes epithelial-mesenchymal transition in breast cancer cells by regulating SPDEF/CDH1 signaling. Mol Carcinog. 2019 Jul;58(7):1314–23.

21. Gong B, Li Y, Cheng Z, Wang P, Luo L, Huang H, et al. GRIK3: A novel oncogenic protein related to tumor TNM stage, lymph node metastasis, and poor prognosis of GC. Tumour Biol J Int Soc Oncodevelopmental Biol Med. 2017 Jun;39(6):1010428317704364.

22. Wen X, Han M, Hosoya M, Toshima R, Onishi M, Fujii T, et al. Identification of BRAF Inhibitor Resistance-associated lncRNAs Using Genome-scale CRISPR-Cas9 Transcriptional Activation Screening. Anticancer Res. 2024 Jun;44(6):2349–58.

23. Hartman ML, Czyz M. MITF in melanoma: mechanisms behind its expression and activity. Cell Mol Life Sci CMLS. 2015 Apr;72(7):1249–60.

24. Ruan X, Lai C, Li L, Wang B, Lu X, Zhang D, et al. Integrative analysis of single-cell and bulk multi-omics data to reveal subtype-specific characteristics and therapeutic strategies in clear cell renal cell carcinoma patients. J Cancer. 2024 Oct 18;15(19):6420.

25. Onieva JL, Xiao Q, Berciano-Guerrero MÁ, Laborda-Illanes A, Andrea C de, Chaves P, et al. High IGKC-Expressing Intratumoral Plasma Cells Predict Response to Immune Checkpoint Blockade. Int J Mol Sci. 2022 Aug 15;23(16):9124.

26. Crescioli S, Correa I, Ng J, Willsmore ZN, Laddach R, Chenoweth A, et al. B cell profiles, antibody repertoire and reactivity reveal dysregulated responses with autoimmune features in melanoma. Nat Commun. 2023 Jun 8;14(1):3378.

27. Flippot R, Teixeira M, Rey-Cardenas M, Carril-Ajuria L, Rainho L, Naoun N, et al. B cells and the coordination of immune checkpoint inhibitor response in patients with solid tumors. J Immunother Cancer. 2024 Apr 1;12(4):e008636.

28. Zhao K, Wang D, Zhao X, Wang C, Gao Y, Liu K, et al. WDR63 inhibits Arp2/3-dependent actin polymerization and mediates the function of p53 in suppressing metastasis. EMBO Rep. 2020 Apr 3;21(4):e49269.

29. Rotty JD, Wu C, Bear JE. New insights into the regulation and cellular functions of the ARP2/3 complex. Nat Rev Mol Cell Biol. 2013 Jan;14(1):7–12.

30. Gu X, Li D, Wu P, Zhang C, Cui X, Shang D, et al. Revisiting the CXCL13/CXCR5 axis in the tumor microenvironment in the era of single-cell omics: Implications for immunotherapy. Cancer Lett. 2024 Nov 28;605:217278.

31. Workel HH, Lubbers JM, Arnold R, Prins TM, van der Vlies P, de Lange K, et al. A Transcriptionally Distinct CXCL13+CD103+CD8+ T-cell Population Is Associated with B-cell Recruitment and Neoantigen Load in Human Cancer. Cancer Immunol Res. 2019 May 1;7(5):784–96.

32. Maibach F, Sadozai H, Seyed Jafari SM, Hunger RE, Schenk M. Tumor-Infiltrating Lymphocytes and Their Prognostic Value in Cutaneous Melanoma. Front Immunol. 2020;11:2105.

33. Yang M, Lu J, Zhang G, Wang Y, He M, Xu Q, et al. CXCL13 shapes immunoactive tumor microenvironment and enhances the efficacy of PD-1 checkpoint blockade in high-grade serous ovarian cancer. J Immunother Cancer. 2021 Jan 1;9(1):e001136.

34. Sorin M, Karimi E, Rezanejad M, Yu MW, Desharnais L, McDowell SAC, et al. Single-cell spatial landscape of immunotherapy response reveals mechanisms of CXCL13 enhanced antitumor immunity. J Immunother Cancer. 2023 Feb 1;11(2):e005545.

35. Melnikova VO, Dobroff AS, Zigler M, Villares GJ, Braeuer RR, Wang H, et al. CREB Inhibits AP-2α Expression to Regulate the Malignant Phenotype of Melanoma. PLoS ONE. 2010 Aug 27;5(8):e12452.

36. Wong D, Yin Y. Immune micro-environment analysis and establishment of response prediction model for PD-1 blockade immunotherapy in glioblastoma based on transcriptome deconvolution. J Cancer Res Clin Oncol. 2023 Oct;149(13):11689–703.

37. Cheng ZY, He TT, Gao XM, Zhao Y, Wang J. ZBTB Transcription Factors: Key Regulators of the Development, Differentiation and Effector Function of T Cells. Front Immunol. 2021 Jul 19;12:713294.

38. Hossain SM, Gimenez G, Stockwell PA, Tsai P, Print CG, Rys J, et al. Innate immune checkpoint inhibitor resistance is associated with melanoma sub-types exhibiting invasive and de-differentiated gene expression signatures. Front Immunol [Internet]. 2022 Sep 28 [cited 2025 Mar 4];13. Available from: https://www.frontiersin.org/journals/immunology/articles/10.3389/fimmu.2022.955063/full

39. Liu J, Chen H, Qiao G, Zhang JT, Zhang S, Zhu C, et al. PLEK2 and IFI6, representing mesenchymal and immune-suppressive microenvironment, predicts resistance to neoadjuvant immunotherapy in esophageal squamous cell carcinoma. Cancer Immunol Immunother CII. 2023 Apr;72(4):881–93.

40. Benci JL, Xu B, Qiu Y, Wu TJ, Dada H, Twyman-Saint Victor C, et al. Tumor Interferon Signaling Regulates a Multigenic Resistance Program to Immune Checkpoint Blockade. Cell. 2016 Dec 1;167(6):1540–1554.e12.

41. Yang J, Qi M, Fei X, Wang X, Wang K. Long non-coding RNA XIST: a novel oncogene in multiple cancers. Mol Med. 2021 Dec 20;27(1):159.

42. Moutafi MK, Bates KM, Aung TN, Milian RG, Xirou V, Vathiotis IA, et al. High-throughput transcriptome profiling indicates ribosomal RNAs to be associated with resistance to immunotherapy in non-small cell lung cancer (NSCLC). J Immunother Cancer [Internet]. 2024 Jun 10 [cited 2025 Jul 17];12(6). Available from: https://jitc.bmj.com/content/12/6/e009039

43. Tay RE, Richardson EK, Toh HC. Revisiting the role of CD4+ T cells in cancer immunotherapy-new insights into old paradigms. Cancer Gene Ther. 2021 Feb;28(1–2):5–17.

44. Vanmeerbeek I, Borras DM, Sprooten J, Bechter O, Tejpar S, Garg AD. Early memory differentiation and cell death resistance in T cells predicts melanoma response to sequential anti-CTLA4 and anti-PD1 immunotherapy. Genes Immun. 2021 Jun;22(2):108–19.

45. Flach H, Rosenbaum M, Duchniewicz M, Kim S, Zhang SL, Cahalan MD, et al. Mzb1 Protein Regulates Calcium Homeostasis, Antibody Secretion, and Integrin Activation in Innate-like B Cells. Immunity. 2010 Nov 24;33(5):723–35.

46. Song L, Ouyang Z, Cohen D, Cao Y, Altreuter J, Bai G, et al. Comprehensive Characterizations of Immune Receptor Repertoire in Tumors and Cancer Immunotherapy Studies. Cancer Immunol Res. 2022 Jul 1;10(7):788–99.

47. Zhu M, Zhou G, Chang F, Liu J. MZB1 regulates the immune microenvironment and inhibits ovarian cancer cell migration. Open Med. 2025 May 13;20(1):20251174.

48. Cabrita R, Lauss M, Sanna A, Donia M, Skaarup Larsen M, Mitra S, et al. Tertiary lymphoid structures improve immunotherapy and survival in melanoma. Nature. 2020 Jan;577(7791):561–5.

49. Helmink BA, Reddy SM, Gao J, Zhang S, Basar R, Thakur R, et al. B cells and tertiary lymphoid structures promote immunotherapy response. Nature. 2020 Jan;577(7791):549–55.

50. Chang TG, Spathis A, Schäffer AA, Gavrielatou N, Kuo F, Jia D, et al. Tumor and blood B-cell abundance outperforms established immune checkpoint blockade response prediction signatures in head and neck cancer. Ann Oncol. 2025 Mar 1;36(3):309–20.

51. Jiang P, Gu S, Pan D, Fu J, Sahu A, Hu X, et al. Signatures of T cell dysfunction and exclusion predict cancer immunotherapy response. Nat Med. 2018 Oct;24(10):1550–8.

52. Subrahmanyam PB, Dong Z, Gusenleitner D, Giobbie-Hurder A, Severgnini M, Zhou J, et al. Distinct predictive biomarker candidates for response to anA-CTLA-4 and anA-PD-1 immunotherapy in melanoma patients. J Immunother Cancer. 2018 Mar 6;6(1):18.

53. Kerle IA, Gross T, Kögler A, Arnold JS, Werner M, Eckardt JN, et al. Translational and clinical comparison of whole genome and transcriptome to panel sequencing in precision oncology. Npj Precis Oncol. 2025 Jan 10;9(1):1–10.

54. Borch TH, Andersen R, Ellebaek E, Met Ö, Donia M, Svane IM. Future role for adoptive T-cell therapy in checkpoint inhibitor-resistant metastatic melanoma. J Immunother Cancer. 2020 Jul;8(2):e000668.

