## Supplementary Material for "Integrating tumor and immune cell transcriptomics to predict immune checkpoint inhibitor primary resistance in metastatic cutaneous melanoma"

**Supplementary Figure**

**Supplementary figure S1. Survival analysis based on primary resistance in the Discovery cohort.** Kaplan-Meier curves illustrating the (A) Progression-free Survival (PFS) and the (B) Overall Survival (OS) for the resistant and non-resistant patients.

**Supplementary figure S2. Heatmap of key marker expression for the 12 imnune cell types from scRNA-seq data.** Heatmap showing the expression of the top 10 markers for each of the 12 immune cell types identified in the single-cell analysis through sc-type.

**Supplementary figure S3. Ridge plot of upregulated DE genes in scRNA-seq across primary resistance and immune cell types**. Ridge plot showing the expression of differentially upregulated genes at the single-cell level, identified in the scRNA-seq of PBMCs, comparing non-resistant and resistant patients with primary resistance. The data is segregated by primary response status and cell type.

**Supplementary figure S4. Ridge plot of downregulated DE genes in scRNA-seq across primary resistance and immune cell types**. Ridge plot showing the expression of differentially downregulated genes at the single-cell level, identified in the scRNA-seq of PBMCs, comparing non-resistant and resistant patients with primary resistance. The data is segregated by primary response status and cell type.

**Supplementary figure S5. Heatmap of key marker expression of Naive CD4$^{+}$ T cells, Memory CD4$^{+}$ T cells, Plasma cells, and Pre-B cells from scRNA-seq data.** Heatmap showing the expression of *TSHZ2, TRABD2A, TESPA1, TOMM7, XBP1, MZB1, BLK, MS4A1, EBF1*, and *PAX5* across the different immune cell types.

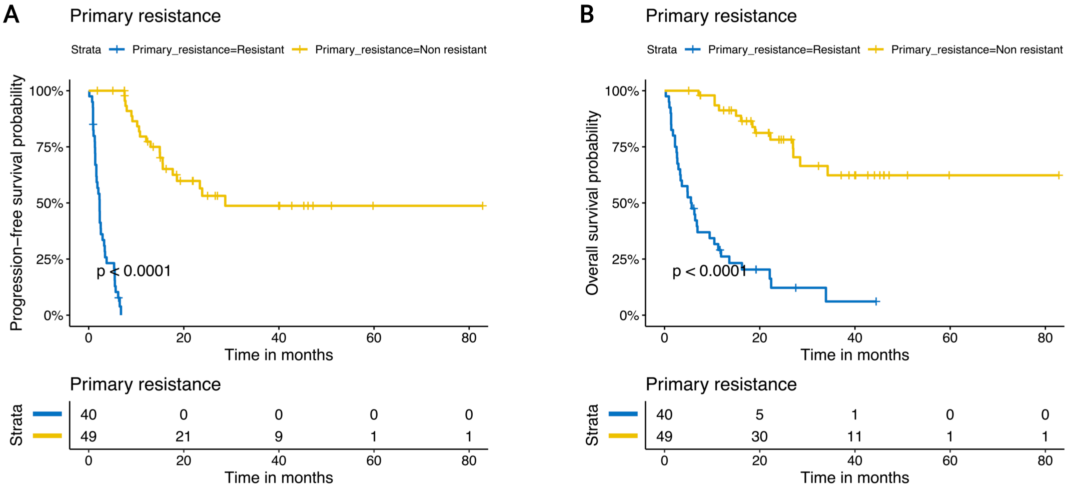


**Supplementary figure S1. Survival analysis based on primary resistance in the Discovery cohort.** Kaplan-Meier curves illustrating the (A) Progression-free Survival (PFS) and the (B) Overall Survival (OS) for the resistant and non-resistant patients.

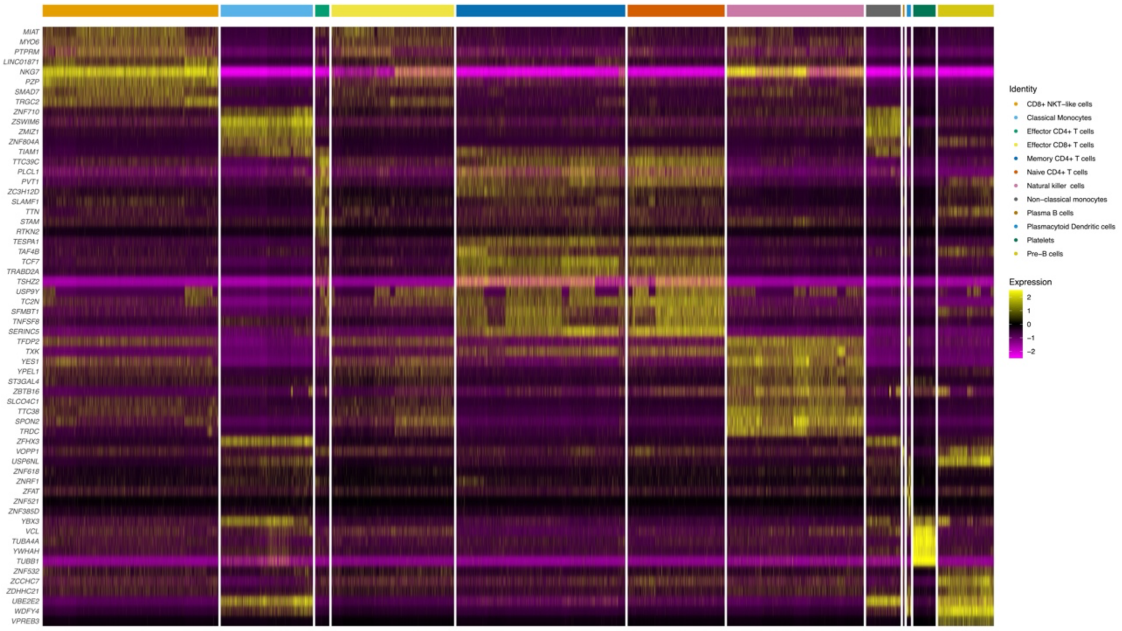


**Supplementary figure S2. Heatmap of key marker expression for the 12 imnune cell types from scRNA-seq data.** Heatmap showing the expression of the top 10 markers for each of the 12 immune cell types identified in the single-cell analysis through sc-type.

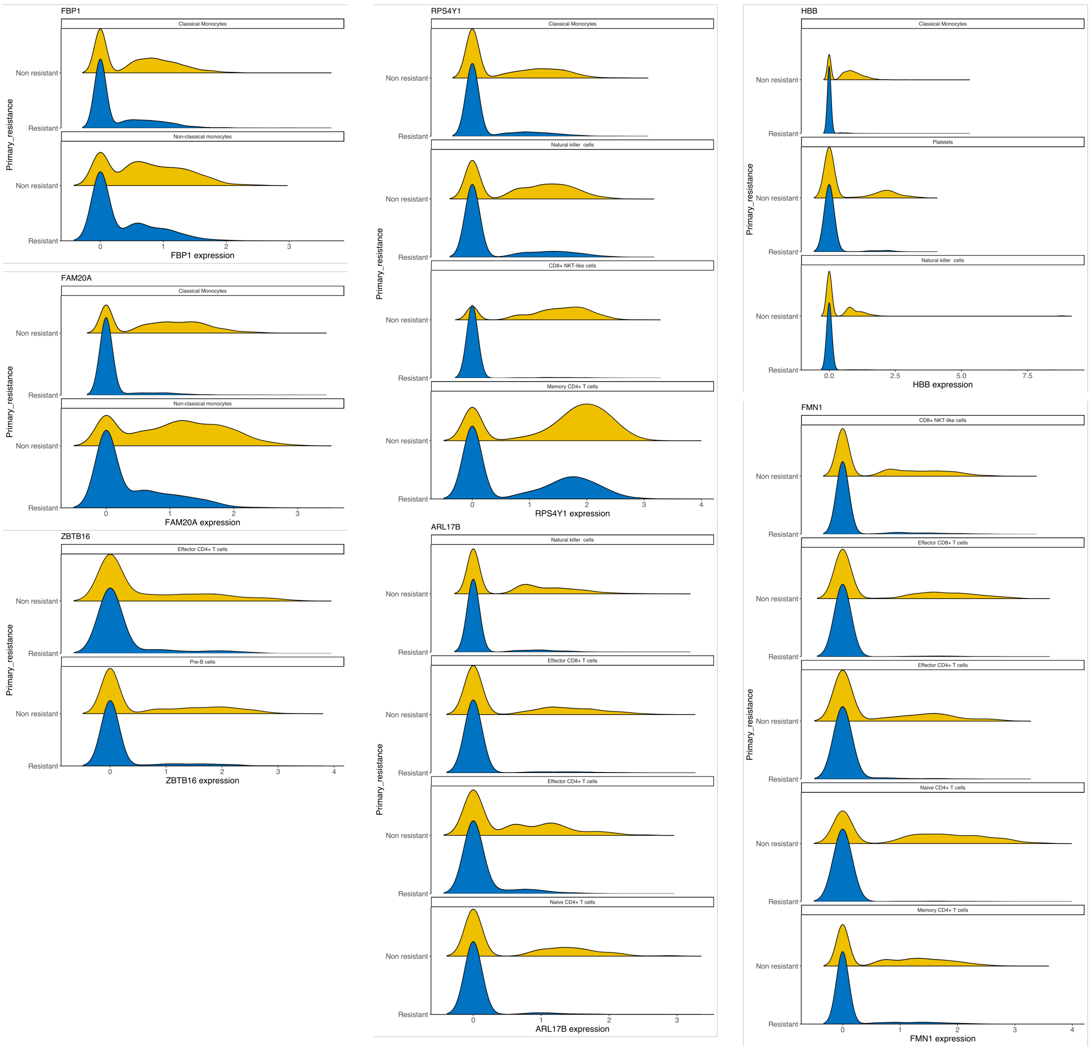


**Supplementary figure S3. Ridge plot of upregulated DE genes in scRNA-seq across primary resistance and immune cell types**. Ridge plot showing the expression of differentially upregulated genes at the single-cell level, identified in the scRNA-seq of PBMCs, comparing non-resistant and resistant patients with primary resistance. The data is segregated by primary response status and cell type.

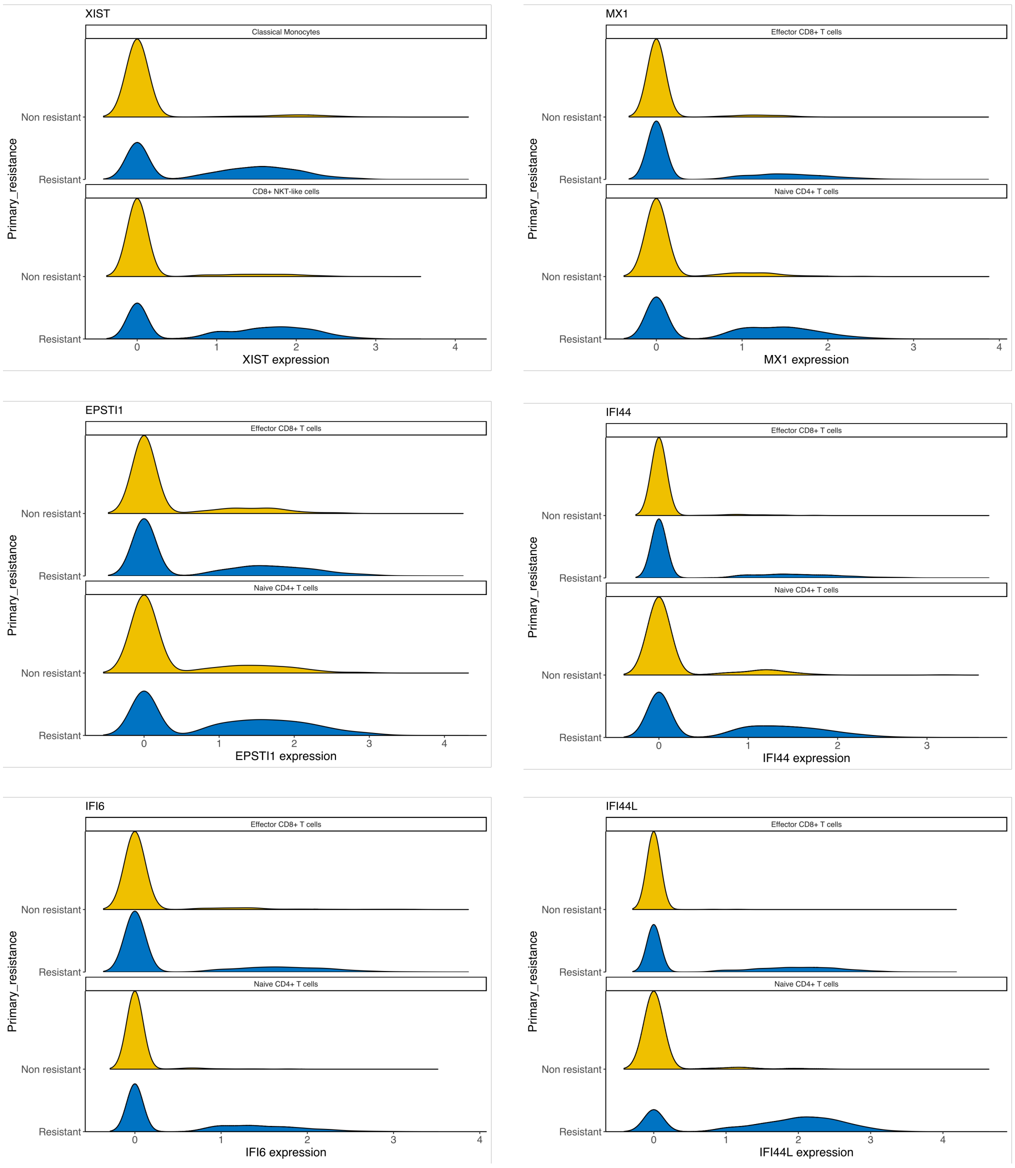


**Supplementary figure S4. Ridge plot of downregulated DE genes in scRNA-seq across primary resistance and immune cell types**. Ridge plot showing the expression of differentially downregulated genes at the single-cell level, identified in the scRNA-seq of PBMCs, comparing non-resistant and resistant patients with primary resistance. The data is segregated by primary response status and cell type.

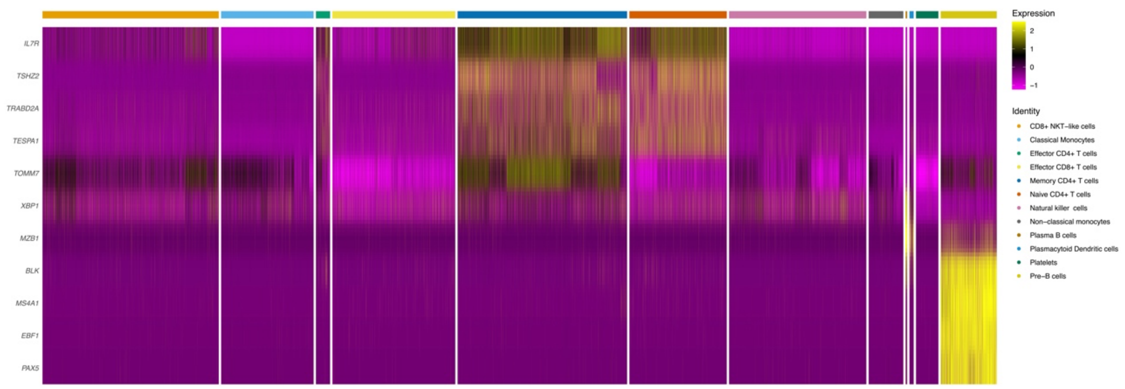


**Supplementary figure S5. Heatmap of key marker expression of Naive CD4$^+$ T cells, Memory CD4$^+$ T cells, Plasma cells, and Pre-B cells from scRNA-seq data.** Heatmap showing the expression of *TSHZ2, TRABD2A, TESPA1, TOMM7, XBP1, MZB1, BLK, MS4A1, EBF1*, and *PAX5* across the different immune cell types.

**Supplementary Table**

**Supplementary table S1.** Clinicopathological characteristics of the Discovery cohort, Single-cell cohort and Flow cytometry cohort.

**Supplementary table S2.** Differentially expressed genes identified from bulk RNA-seq analysis of resistant vs non-resistant patients in the Discovery cohort

**Supplementary table S3.** DE analysis between resistant and non resistant patients from the Single-cell Cohort across cell-types.

**Supplementary table S4.** Top marker genes identified from single-cell RNA-seq analysis for each of the 12 immune cell types inferred using ScType.

**Supplementary table S5.** Deconvolution scores estimating the relative abundance of each immune cell type for each bulk RNA-seq sample from the Discovery cohort.

**Supplementary table S6.** Deconvolution scores estimating the relative abundance of each immune cell type for each bulk RNA-seq sample from the study cohort.

**Supplementary table S7**. Sequencing coverage and quality statistics. Reference Genome GRCh38.

**Supplementary table S8**. Summary of genes of interest.
